# Differential expression of *NEAT1* in the corneal endothelium increases susceptibility to oxidative stress in Fuchs Endothelial Corneal Dystrophy

**DOI:** 10.64898/2026.08.31.748332

**Authors:** Narisa Dhupar, Judy Yan, Ness Little, Stephan Ong Tone

## Abstract

Fuchs endothelial corneal dystrophy (FECD) is a disease of the corneal endothelium (CE) characterized by the loss of corneal endothelial cells (CECs) and guttae formation, ultimately resulting in corneal edema and vision loss. FECD primarily affects the central CE while sparing the peripheral CE, however the underlying mechanism contributing to the spatial differences remain unknown. Oxidative stress has been increasingly recognized as a key contributor to the pathogenesis of FECD, with CECs being particularly susceptible to damage from reactive oxygen species (ROS), high metabolic activity and ultraviolet induced-DNA damage. The non-proliferative nature of CECs, along with the accumulation of oxidative damage can ultimately lead to CEC loss, a key feature of FECD. In this study, we induced oxidative stress with hydrogen peroxide (H2O_2_) on ex vivo corneal specimens and observe increased cell death in the central region compared to the peripheral CE. To investigate these underlying differences, we performed bulk RNA sequencing (RNA-seq) on the central and peripheral regions of CE from FECD and normal cadaveric donors. Pathway analysis identified an enrichment of genes involved in collagen and extracellular matrix between the central and peripheral regions of CE in both normal and FECD, as well as between normal and FECD CE. Intriguingly, we identified the long non-coding RNA (lncRNA), *NEAT1* as a top differentially expressed gene, with reduced expression in the central CE compared to the peripheral CE and lower expression in FECD compared with normal CE. Using corneal endothelial cell lines and ex vivo specimens from FECD patients and normal cadavers, we found decreased *NEAT1* expression levels in FECD and increased susceptibility to H_2_O_2_–induced oxidative stress. We observed that *NEAT1* knockdown in normal and FECD cells exacerbated H_2_O_2_-mediated oxidative stress, and that *NEAT1* overexpression protected FECD cells. We report in this study, a novel insight in the spatial differences in gene expression in the CE and identify reduced expression of *NEAT1* in the central CE as a potential contributor to oxidative stress-related cell death in FECD. These findings provide novel insight into FECD pathogenesis and why FECD pathology preferentially affects the central CE. Antioxidants targeting *NEAT1* signaling could be developed into novel therapeutics aimed at preventing FECD pathogenesis.

## Introduction

Fuchs endothelial corneal dystrophy (FECD) is a progressive, age-related disease with a female predominance that is a leading cause of corneal dysfunction and is the top indication for corneal transplantation worldwide [1,2]. FECD is characterized by progressive corneal endothelial cell (CEC) loss, the formation of corneal guttae, and abnormal Descemet’s membrane (DM) thickening [3–5]. Clinically, FECD presents with blurred vision and glare, and in advanced cases, corneal edema and vision loss [6]. Although the pathogenesis of FECD remains to be completely elucidated, both genetic and environmental factors, such as ultraviolet light and oxidative stress have been reported to contribute to its onset [7,8].

The corneal endothelium (CE) is composed of a single monolayer of hexagonal cells situated on DM on the posterior side of the cornea [9]. Its primary function is to preserve corneal deturgescence by actively pumping out fluid from the stroma to maintain corneal clarity [10]. At birth, CE density is approximately 4,000 cells/mm^2^ and subsequently decreases with age averaging around 2000 cells/mm^2^ in healthy adults [11,12]. CECs are quiescent in nature and are arrested in the G1 phase of the cell cycle [13]. Due to its limited proliferative capacity, the CE relies substantially on cell enlargement and cell migration to facilitate wound healing to recapitulate a monolayer after cell loss [13–16]. Under pathological conditions, such as FECD, loss of CECs can lead to a disruption of the CE layer and compromise function.

The progression of FECD is well documented, beginning with early CEC loss and guttae formation, typically confined to the central cornea, while the periphery remains unaffected. As FECD progresses, there is further cell loss that preferentially affects the central CE leading to CEC apoptosis and decrease endothelial cell density [8,17]. Accumulating evidence indicates the CE is not a uniform cell population but rather consists of subpopulations of CECs with unique properties. A surgical technique for FECD, termed Descemet Stripping Only (DSO), has been developed whereby the central diseased CE and DM is excised, and relies on peripheral CEC migration to the center to regenerate a functional endothelium. DSO success is based on the spatial differences in FECD pathology, where the center CE is preferentially affected, and the peripheral CECs can migrate centrally. Moreover, the identification of progenitor-like cells in the inner transition zone at the posterior limbus of the corneal periphery further indicates the CE is comprised of a heterogenous cell population [18]. Recently, single cell RNA sequencing (RNA-seq) identified the existence of 4 distinct sub population of CECs by unique genetic signatures [19]. Given the heterogenous nature of the CE and the pathological changes observed in FECD that primarily affects the central CE while sparing the periphery, the underlying cause of this spatial distribution pattern during FECD pathology remains unknown and represents a significant clinical gap yet to be addressed.

Oxidative stress arises from an imbalance characterized by either a reduction in antioxidant defenses, an accumulation of reactive oxygen species (ROS), or both. In response, cells modulate gene expression—activating or repressing specific genes—to restore redox homeostasis. ROS and H_2_O_2_ signaling can damage macromolecules, including lipids and proteins, and lead to DNA damage and apoptosis [20,21]. As the eye is continuously exposed to ultraviolet (UV) radiation, cellular damage through oxidative stress has been linked to a number of non-cancer ocular pathologies, including cataract, glaucoma and age-related macular degeneration [21]. In the cornea, continuous ultraviolet exposure results in elevated oxidative stress within CECs, with evidence indicating a greater burden in the central CE compared to the periphery [22]. Evidence shows UV light and oxidative stress can induce cellular senescence, DNA damage, cytotoxicity and apoptosis in CECs [23–29]. In mouse cornea, UV-A irradiation induced ROS production, CEC loss and greater mitochondria DNA and nuclear DNA damage [30]. Substantial evidence indicates that oxidative stress is a critical contributor to FECD pathogenesis, as reflected by reduced expression of antioxidant genes in the CE and markedly elevated oxidative DNA damage and apoptosis in FECD specimens compared to normal controls [25–27]. Wang *et al.* [19] identified Nuclear Paraspeckle Assembly Transcript 1, *NEAT1*, an antioxidant gene, highly expressed in a CEC subtype and was significantly decreased in FECD samples [19]. In a non-genetic FECD animal model induced by UV-A, as well as CEC lines, CEC death was exacerbated by *NEAT1* knockdown, and attenuated by *NEAT1* overexpression, indicating *NEAT1* as an important antioxidant in the CE [19]. *NEAT1* is a regulatory long non-coding RNA (lncRNA) found to localize with nuclear bodies, specifically paraspeckles [19,31,32]. To date, two isoforms have been reported, a 3,700bp (*NEAT1* short) isoform that completely overlaps with the 5’ end of the 23,000bp (*NEAT1* long) isoform [32]. *NEAT1* has been reported to be essential for paraspeckle formation and has furthermore been implicated as a regulatory factor in the DNA damage response [31,33–35]. Pathologically, *NEAT1* has been associated with a variety of cancers [36–38] and confers neuroprotective effects from oxidative stress in Parkinson’s disease [39], as well as inhibiting oxidative stress induced vascular endothelial cell damage [40].

In this study, we observed increased cell death in the central region of ex vivo corneal specimens following H_2_O_2_-induced oxidative stress. RNA-seq on spatially dissected central and peripheral regions of CE from normal and FECD specimens identified *NEAT1* as a top differentially expressed gene. Using ex vivo specimens and cell lines derived from normal cadaveric donors and FECD patients we observed lower endogenous *NEAT1* levels and increased sensitivity to H_2_O_2_-induced oxidative stress in FECD. *NEAT1* knockdown further exacerbated oxidative stress induced cell death in normal and FECD cells, whereas *NEAT1* over expression conferred protection against H_2_O_2_-mediated cell death in FECD cells. To our knowledge, this is the first evidence demonstrating the central CE is more susceptible to oxidative stress and this susceptibility may in part be attributed to reduced levels of the antioxidant gene, *NEAT1*. These findings offer novel insight into the spatial differences in FECD pathology.

## Materials and Methods

### Human tissues

The study was carried out according to the tenets of The Declaration of Helsinki and was approved by the Sunnybrook Health Sciences Centre Research Ethics Board (REB#5070 and REB#5187). Normal and FECD tissues from cadaveric donors were provided by The Eye Bank of Canada and Eversight USA and stored in Optisol-GS (Bausch & Lomb) at 4°C until use. FECD diagnosis was made based on specular microscopy examination of guttae in the corneal endothelium or by medical history during time of tissue collection (Figure S1). FECD surgical specimens were collected from patients undergoing surgery for transplant following informed consent. See Supplementary Table 1 for patient and donor characteristics.

### Cell lines

Immortalized FECD cell lines were generated as previously described [41]. FECD-SVF5-54F and FECD-SVF7-74F was isolated from a 54-year-old and 74-year-old female, respectively, undergoing endothelial keratoplasty. HCEC-SVN1-67F was isolated from a 67-year-old female deceased donor. CECs were isolated and SV-40 immortalized to generate cell lines [41]. HCEC-SVN4-68F (68-year-old female) and FECD-SVF6-61M (61-year-old male) were graciously provided by Dr. Ula Jurkunas at The Schepens Eye Research Institute in Boston, Massachusetts. Cells were cultured in Chen’s Media, containing Opti-MEM media (ThermoFisher Scientific, Waltham, MA) supplemented with 200 mg/L CaCl_2_ (Millipore Sigma, Oakville, Ontario), 0.08% chondroitin sulfate (Millipore Sigma, Oakville, Ontario), 50 μg/mL gentamicin (ThermoFisher Scientific, Waltham, MA), 1 × antibiotic/antimycotic (Wisent, St. Bruno, QC), 66 μg/mL bovine pituitary extract (Gemini, West Sacramento, CA), 5 ng/mL epidermal growth factor (Millipore Sigma, Oakville, Ontario) and 8% fetal bovine serum (ThermoFisher Scientific, Waltham, MA). Cell culture flasks or plates were pre-coated with undiluted fibronectin coating mix (AthenaES, Baltimore, MD) before seeding and incubated at 37°C with 5% CO_2_.

### Tissue Processing and RNA isolation

Descemet’s membrane and CE were collected from normal (n=3) and FECD cadaveric donors (n=3). The corneoscleral tissue was placed on an 8-mm diameter Barron corneal vacuum donor punch. Approximately 50% of the CE and DM was gently peeled using fine non-toothed forceps before being placed back into position and trephined to separate out the 8-mm central CE and remaining peripheral CE. Total RNA was isolated from each spatial domain using the PureLink RNA Micro Kit (ThermoFisher Scientific, Waltham, MA) according to the manufacturer’s protocol. Samples were sent to the Genomics Core Facility at Princess Margaret Cancer Centre in Toronto, Canada and the quantity of total RNA were determined using an Agilent 2100 Bioanalyzer with an RNA 6000 Pico Kit (Agilent Technologies, Santa Clara, CA). The quality of total RNA was assessed by RNA integrity number (RIN) using the Agilent 2100 Expert Software (Agilent Technologies).

### RNA-Seq library preparation and data processing

The RNA-seq libraries for next-generation sequencing (NGS) were generated with Illumina Stranded Total RNA Ribo Zero Plus kit according to the manufacturer’s instructions. Sequencing was carried out with the NovaSeq 6000 using a 100-cycle paired read protocol and multiplexing to obtain ∼40 million reads/sample, and data was analyzed using bcl-convert Version 4.1.5 to generate differentially expressed genes (DEGs) and FASTQ files.

The read quality on the raw sequencing data was checked using FASTQC v.0.11.5.[42]. The raw sequencing data were aligned to the human genome (GRCh38, Homo_sapiens.GRCh38.84.gtf) using the HISAT2 v2.2.1.[43]. The accessory software tool for the alignment stage includes Samtools v1.17.[44]. Transcript assembly was completed using StringTie v2.1.4.[45]. HTSeq (v0.11.0) toolkit was utilized to access the htseq-count tool which pre-processed the RNA-seq alignments and generated read counts. Alignment quality control was completed using Picard v2.10.9 (“Picard Toolkit”) and MultiQC v1.7.[46]. The Ballgown R package v2.36.0 was utilized for downstream analysis [47]. Low-abundance genes were removed by calculating the variance per transcript. Differential expression analyses were conducted using the stattest() function on the transcripts and the genes, separately. Figures were created using ggplot2 v3.5.1 and EnhancedVolcano R packages v1.22.0 [48]. The following auxiliary packages were used for the analysis: tidyverse v2.0.0 [49], genefilter v1.82.1, dendextend v1.17.1 [50] and ggdendro v0.2.0. Genes with fold change > 1.5 and < -1.5 and p-value < 0.05 were selected as differentially expressed genes (DEGs). Gene ontology (GO) and pathway enrichment analysis was carried out using g:Profiler [51].

### RNAscope

Corneal cadaveric donor tissues were fixed in 4% paraformaldehyde (PFA) for 24 hours at 4°C and cryo-embedded in Tissue-Tek optimal cutting temperature (OCT) compound (Fisher Scientific, Pittsburgh, PA). Tissues were sectioned at 10 μm thickness and probed for *NEAT1* using RNAscope Multiplex Fluorescent V2 Assay Kit (ACD Bio, Newark, CA) according to manufacturer’s protocol. Hs-NEAT1-short-C3 and Hs-NEAT1-C1 probes were used to detect the short and long isoform of *NEAT1,* respectively. Samples were counterstained with DAPI (ACD Bio, Newark, CA) and mounted with ProLong Diamond Antifade Mountant (Thermo Fisher Scientific, Waltham, MA). Images were captured using a Nikon A1 laser scanning confocal microscope (Nikon Instruments Inc., Melville, NY). The central 8-mm corresponding to the central CE was approximated, and images were acquired for the central CE and the peripheral CE.

### Cell Viability Assay

Descemet’s membrane and the CE was stripped from the corneoscleral rims and placed in 1.9 mM hydrogen peroxide (H_2_O_2_) solution in Chen’s media (Sigma Aldrich, St. Louis, MO) to induce oxidative stress. Tissues were incubated at 37°C for 2 hours. LIVE/DEAD® Viability/Cytotoxicity Assay Kit (Molecular Probes, Eugene, OR) at 3.12 µM calcein AM (CAM), 4 µM Ethidium homodimer-1 (EthD-1), and 4 µM Hoescht-33342 (Thermo Fisher Scientific) was reconstituted in 1.7% hyaluronic acid (HA; Sigma-Aldrich, St. Louis, MO) in Hank’s Balanced Salt Solution (HBSS). Using a no-touch technique, tissues were gently unfolded on to a glass slide and mounted with a coverslip prior to imaging. For cells, HCEC-SVN4-68F and FECD-SVF7-74F were seeded at a density of 100,000 cells per well in 24 well plates pre-coated with undiluted fibronectin coating mix overnight. H_2_O_2_ diluted in Chen’s media was added to cells and incubated at 37°C for 2 hours. LIVE/DEAD® Viability/Cytotoxicity Assay Kit (Molecular Probes, Eugene, OR) at 3.12 µM calcein AM (CAM), 4 µM Ethidium homodimer-1 (EthD-1), and 4 µM Hoescht-33342 in 1xPBS was added to cells and imaged. All images were acquired with a Leica S DMi8 inverted fluorescence microscope. For tissues, the approximated central 8 mm region was measured using the Leica Application Suite X (Version 3.8.1.26810), and images were acquired for the central CE and the peripheral CE. Cell viability quantification was performed in ImageJ, as previously reported [52].

### Quantitative real-time PCR

Total RNA was extracted using the PureLink RNA Mini Kit (Thermo Fisher Scientific, Waltham, MA) or PureLink RNA Micro Kit according to the manufacturer’s protocol. Reverse transcription was carried out using Superscript IV (ThermoFisher Scientific, Waltham, MA). Quantitative PCR was performed using the StepOnePlus System (Applied Biosystems, Foster City, CA) with Power SYBR Green PCR Master Mix (Applied Biosystems, Foster City, CA) according to the manufacturer’s protocol. Relative gene expression levels were calculated using ΔΔCt. Primers used for qRT-PCR are listed in Supplementary Table 2.

### Lentiviral knockdown and overexpression of NEAT1 in cell lines

Lentiviral vectors pLV[Exp]-EGFP:T2A:Hygro-EFS-hNEAT1 expressing Human NEAT1 short isoform (NR_028272.1) and pLV[shRNA]-EGFP:T2A:Hygro-U6-hNEAT1 targeting NEAT1 (Target sequence 5’ACGCAGCAGATCAGCATCCTT 3’) were constructed by Vector Builder (Chicago, IL). pLV[Exp]-EGFP:T2A:Hygro-EFS-MCS vector without NEAT1 was used as an empty vector control, and pLV[shRNA]-EGFP:T2A:Hygro-U6-ScrambleshRNA (Target Sequence 5’CAACAAGATGAAGAGCACCAA 3’) was used as a control for knockdown. A 2nd generation lentiviral packaging system, psPAX2 and pMD2.G plasmids was transiently co-transfected with the designed lentiviral plasmid with Lipofectamine 3000 (Thermo Fisher Scientific, Waltham, MA) into HEK293T. The virus containing media was collected 48 hours later and centrifuged to remove cell debris. Viral supernatant was filtered through a 0.45μm filter and added to cells for 24 hours. Green fluorescence protein (GFP) expression was visualized for transduction efficiency and selected for stable integration with hygromycin (200 μg/mL, Wisent, St. Bruno, QC). NEAT1 knockdown and overexpression were confirmed using qRT-PCR.

### Statistical Analysis

Statistical analysis was performed using a student *t-*test or either a one-way or two-way analysis of variance (ANOVA) followed by a Tukey post hoc test. A p*-*value <0.05 was considered statistically significant. Values shown in graphs represent mean±SEM.

## Results

### The central CE is more susceptible to H_2_O_2_-mediated cell death

In FECD, disease begins in the center with early guttae formation and corneal endothelial cell (CEC) loss, with dysregulated oxidative stress playing a key role in its pathogenesis [25–27]. To investigate spatial susceptibility to oxidative stress, H_2_O_2_-mediated cell death was examined in cadaveric CE (Figure 1A). The central 8-mm CE exhibited greater sensitivity to H_2_O_2_ induced oxidative stress with 42.4% cell viability in the center compared to 74.1% cell viability in the peripheral CE (Figure 1B,C). The increased susceptibility of oxidative stress in the central region of the CE suggests underlying differences in CECs.

**Figure 1.**
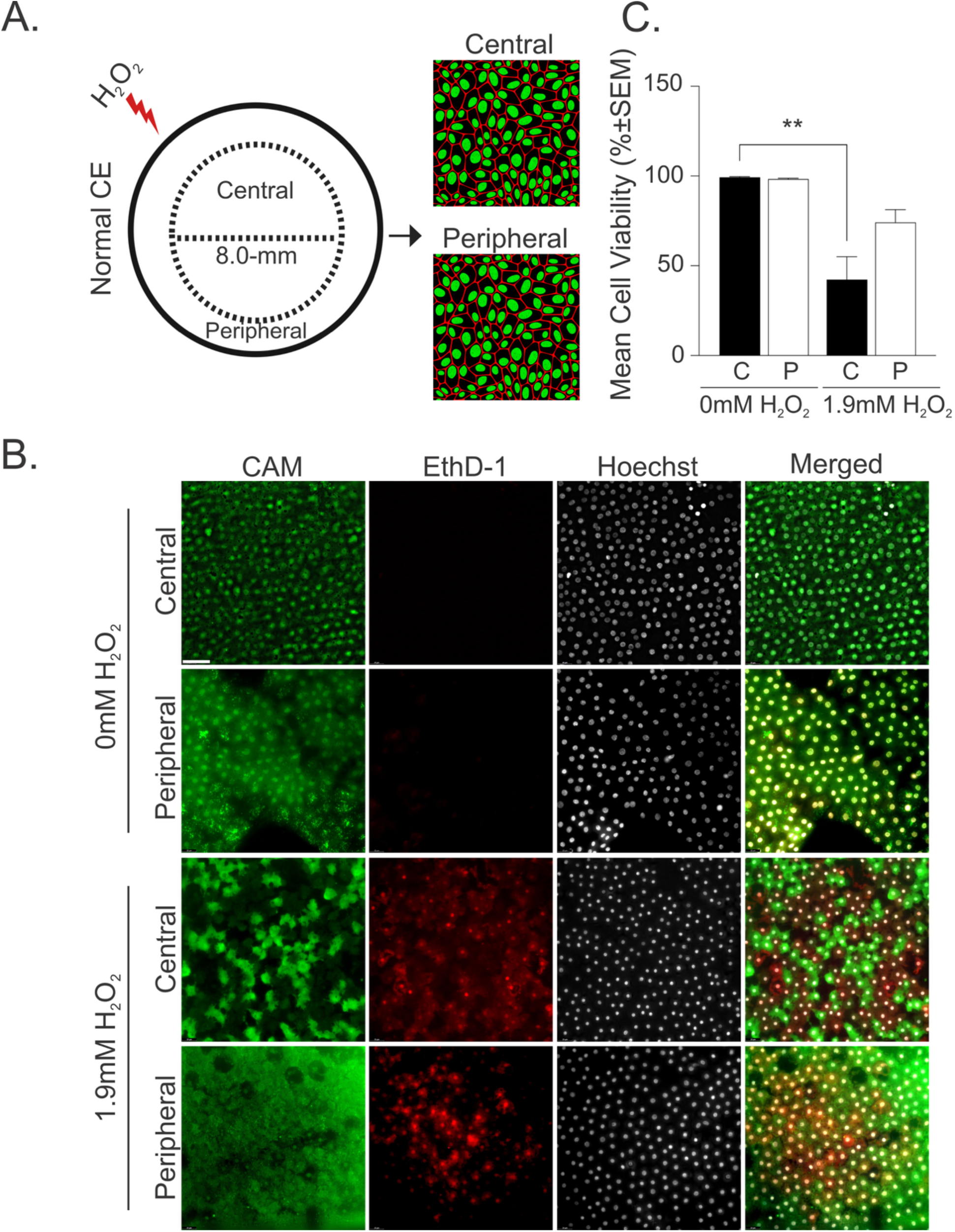
Increased susceptibility to H_2_O_2_-mediated cell death in the central corneal endothelium. **A)** Schematic representation of H_2_O_2_-induced oxidative stress in normal cadaveric tissue followed by imaging of the central and peripheral regions. Approximation of the 8-mm designated central region is shown. **B)** Cell viability staining on normal CE treated with 0 mM and 1.9 mM H_2_O_2_. Representative fluorescence images (scale bar = 50 µm) of central and peripheral CE for live cells (green), dead cells (red) and nuclei staining (grey). **C)** Quantification of cell viability; C=central, P=peripheral. **p-value < 0.01 by two-way ANOVA and Tukey post hoc test. SEM = standard error of the mean.

### Identification of reduced NEAT1 expression in the central CE and FECD

To investigate differentially expressed genes (DEGs) that can attribute to the spatial differences between the central and peripheral CE, we performed bulk RNA sequencing on cadaveric tissues from healthy normal and FECD donors. Equivalent sample areas from the central 8-mm region were dissected and separated from the periphery for both normal and FECD cadaveric donor CE. Multidimensional scaling (MDS) plots reveal more separation within the peripheral CE samples than central CE samples (Figure 2A). We identified 166 DEGs (85 upregulated and 81 downregulated) in normal peripheral compared to central CE (Figure 2B, Supplementary Table 3). Among the top 20 DEGs upregulated (Figure 2C) and downregulated (Figure 2D) in the periphery, the antioxidant gene, *NEAT1* was identified to be significantly upregulated (log_2_FC=1.75, p-value=0.0098) in the peripheral CE compared to the central in normal CE (Figure 2B,C). Gene Ontology (GO) analysis of upregulated DEGs demonstrated enrichment in pathways involved in collagen organization and extracellular matrix modulation (Figure 2E), whereas downregulated DEGs enriched for GO annotations associated with mitochondrial function (Figure 2F). In FECD specimens, FECD peripheral CE overlap closely, with less overlap in central CE samples (Figure 3A). A total of 212 DEGs (186 upregulated and 26 downregulated) were identified between FECD central and peripheral CE (Figure 3B, Supplementary Table 4). In contrast to normal CE, *NEAT1* did not reach statistical significance as an upregulated or downregulated DEG in the periphery (Figure 3C,D, Supplementary Table 4). The upregulated genes in the peripheral FECD CE were enriched for pathways related to ECM organization and remodeling, cell adhesion/migration and developmental processes (Figure 3E). While *NEAT1* was not identified as a DEG between the FECD central and peripheral CE, we observed a trend that its expression was upregulated in the peripheral CE relative to the central (p=0.066, Figure 3F).

**Figure 2.**
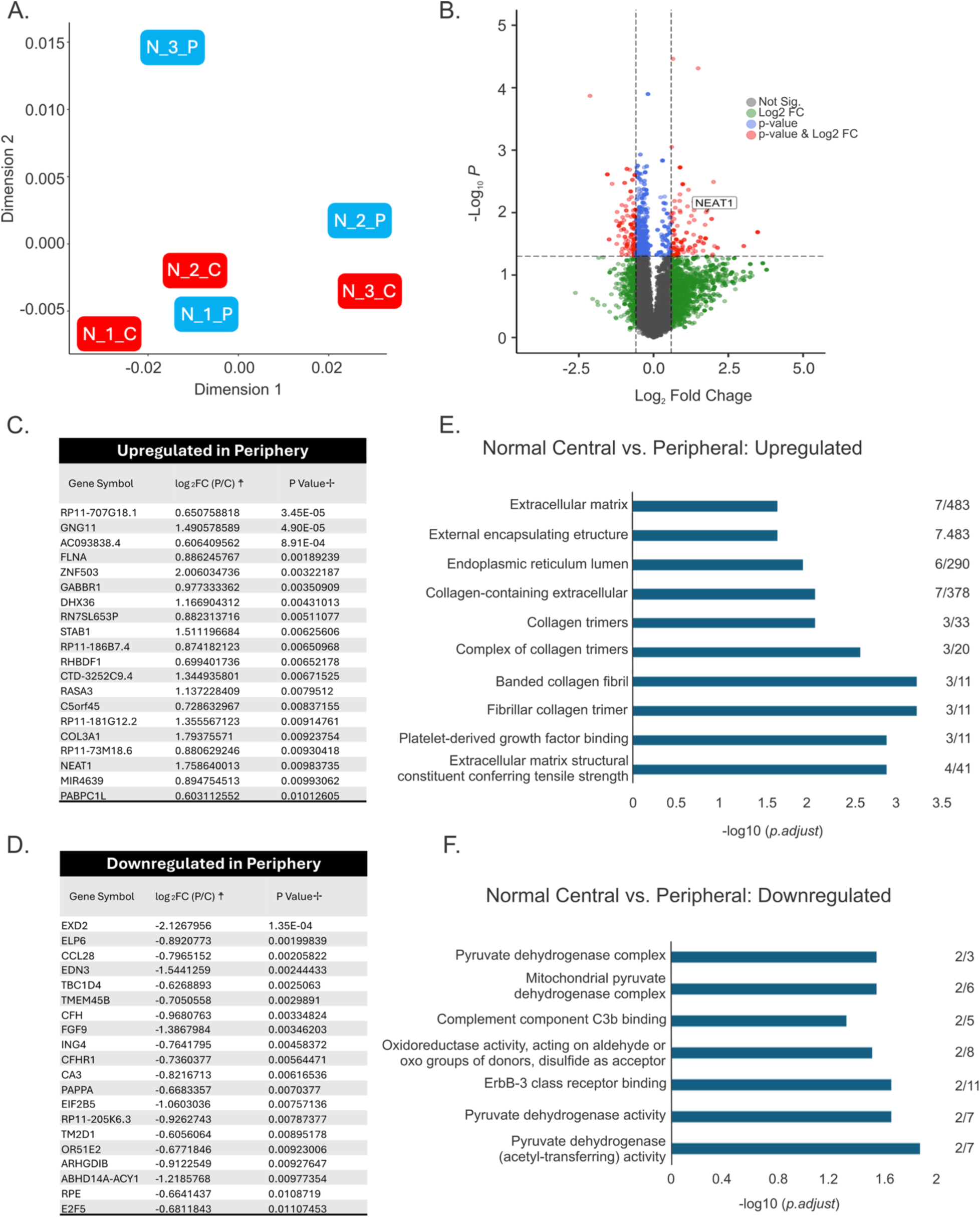
Differentially expressed genes (DEGs) and pathway analysis in normal corneal endothelium between the central and the peripheral spatial domains. A) Multidimensional scaling (MDS) plot for central and peripheral regions for normal CE; C-central, P-peripheral N-normal. **B)** Volcano plot of DEGs (red) showing 85 upregulated and 81 downregulated genes in the periphery compared to central regions at a p-value < 0.05 and Log2 fold change (Log2FC) of > 0.59 and < -0.59 (dashed line). The position of *NEAT1* is marked. Non-significant genes (grey), genes with a p-value < 0.05 and Log2FC < 0.59 and > -0.59 (blue) and genes with a p-value > 0.05 and Log2FC > 0.59 and < -0.59 (green) are indicated. **C)** Top 20 upregulated DEGs in the periphery compared to central CE. **D)** Top 20 downregulated DEGs in the periphery compared to central CE. Pathway enrichment analysis identifying gene ontology (GO) annotations for **E)** upregulated and **F)** downregulated DEGs between periphery compared to central CE. Numbers on the right of each GO annotations is the number of DEGs in the pathway over the total number of genes in the annotation. ☨=Log2 Fold Change (Peripheral/Central). ✢ = p-value by Wald test DESeq2.

**Figure 3.**
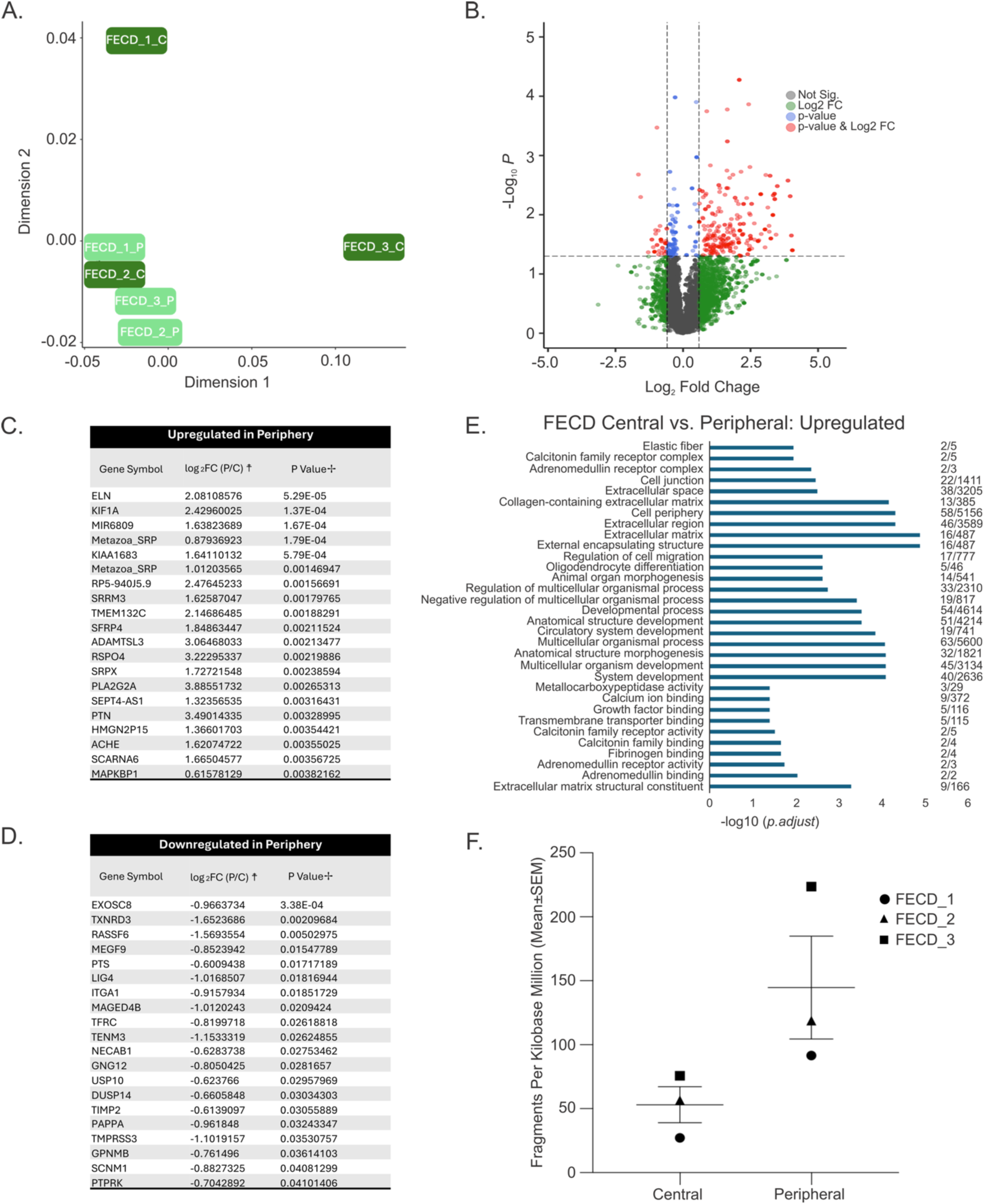
Differentially expressed gene (DEGs) and pathway analysis in FECD corneal endothelium between the central and the peripheral spatial domains. **A)** Multidimensional scaling (MDS) plot for central and peripheral regions for FECD CE; C-central, P-peripheral. **B)** Volcano plot of DEGs (red) showing 186 upregulated and 26 downregulated genes in the peripheral compared to central regions at a p-value < 0.05 and Log2 fold change (Log2FC) of > 0.59 and < -0.59 (dashed line). Non-significant genes (grey), genes with a p-value < 0.05 and Log2FC < 0.59 and > -0.59 (blue) and genes with a p-value > 0.05 and Log2FC > 0.59 and < -0.59 (green) are indicated. **C)** Top 20 upregulated DEGs in the periphery compared to central CE. **D)** Top 20 downregulated DEGs in the periphery compared to central CE. **E)** Pathway enrichment analysis identifying gene ontology (GO) annotations for upregulated DEGs between periphery compared to central CE. Numbers on the right of each GO annotations is the number of DEGs in the pathway over the total number of genes in the annotation. F) *NEAT1* gene expression levels in central and peripheral regions for FECD CE. ☨=Log2 Fold Change (Peripheral/Central). ✢ = p-value by Wald test DESeq2.

To investigate transcriptomic differences associated with FECD, we integrated our spatial transcriptomic datasets and compared gene expression profiles between normal and FECD CE. MDS plots show overlap among normal and FECD samples (Figure 4A) with 367 DEGs (129 upregulated and 238 downregulated) in FECD compared to normal (Figure 4B-D, Supplementary Table 5). GO analysis on genes upregulated in FECD enriched for pathways involved in ECM organization and remodeling, collagen organization and transforming growth factor-β (TGF-β) signaling pathways (Figure 4E). In contrast, genes downregulated in FECD enriched for pathways involved in small molecular binding, metabolic pathways and redox functioning (Figure 4F). While a reduction in *NEAT1* levels has previously been reported in FECD [19], we did not observe a statistically significant difference in *NEAT1* levels in our RNA-seq dataset between normal and FECD CE. However, to further investigate *NEAT1* expression levels in FECD, we examined surgical specimens from patients with a clinical diagnosis of FECD (N=6) to normal CE from cadaveric donors. Since only the central cornea is surgically collected from FECD patients, comparisons were kept to the central region of normal and FECD CE. Using quantitative real-time PCR we observed a significant decrease in *NEAT1* levels in FECD compared to normal (Figure 4G) consistent with previous studies. To further validate the spatial distribution of *NEAT1* expression in the central CE, *NEAT1* levels was assessed using cross sections of normal corneal donor tissues (Figure 5A). *In situ* hybridization confirmed elevated *NEAT1* expression for both the short (total *NEAT1*) (6.1±1.5 vs. 11.5±0.9 dots/cell; Figure 5B,C) and long (6.3±2.8 vs. 9.3±3.1 dots/cell; Figure 5B,D) isoform of *NEAT1* in the peripheral CE compared to the central.

**Figure 4.**
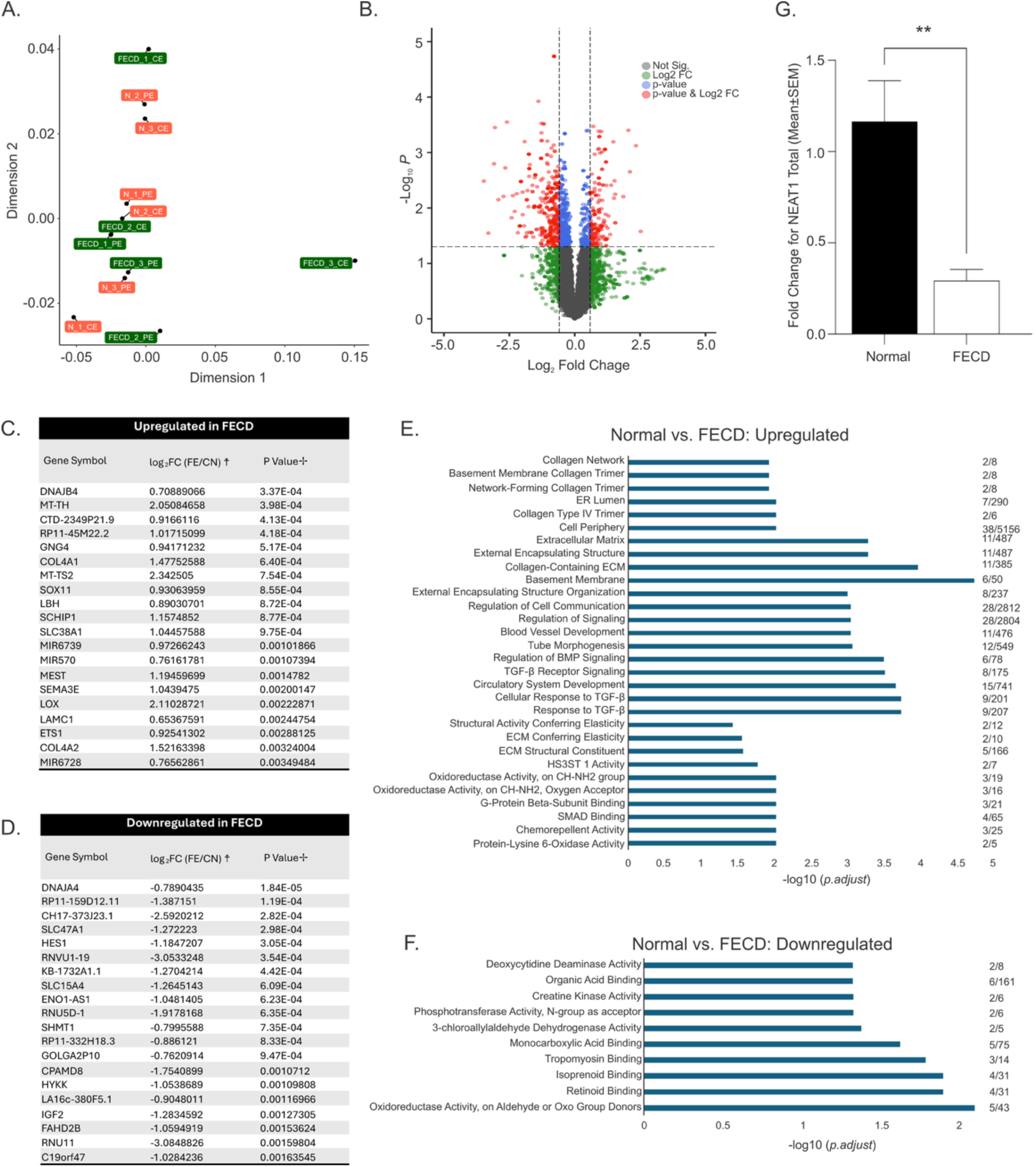
Differentially expressed gene (DEGs) and pathway analysis between normal and FECD corneal endothelium. **A)** Multidimensional scaling (MDS) plot for normal and FECD CE; CE-central endothelium, PE-peripheral endothelium, N-normal **B)** Volcano plot of DEGs (red) showing 129 upregulated and 238 downregulated genes in FECD compared to normal CE at a p-value < 0.05 and Log2 fold change (Log2FC) of > 0.59 and < -0.59 (dashed line). Non-significant genes (grey), genes with a p-value < 0.05 and Log2FC < 0.59 and > -0.59 (blue) and genes with a p-value > 0.05 and Log2FC > 0.59 and < -0.59 (green) are indicated. **C)** Top 20 upregulated DEGs in FECD compared to normal CE. **D)** Top 20 downregulated DEGs in FECD compared to normal CE. Pathway enrichment analysis identifying gene ontology (GO) annotations for **E)** upregulated and **F)** downregulated DEGs between FECD and normal CE. Numbers on the right of each GO annotations is the number of DEGs in the pathway over the total number of genes in the annotation. **G)** *NEAT1* gene expression levels in normal central corneal endothelium and central FECD surgical specimens, **p-value < 0.01 by two-tailed student *t*-test. ☨=Log2 Fold Change (FECD/Normal). ✢ = p-value by Wald test DESeq2.

**Figure 5.**
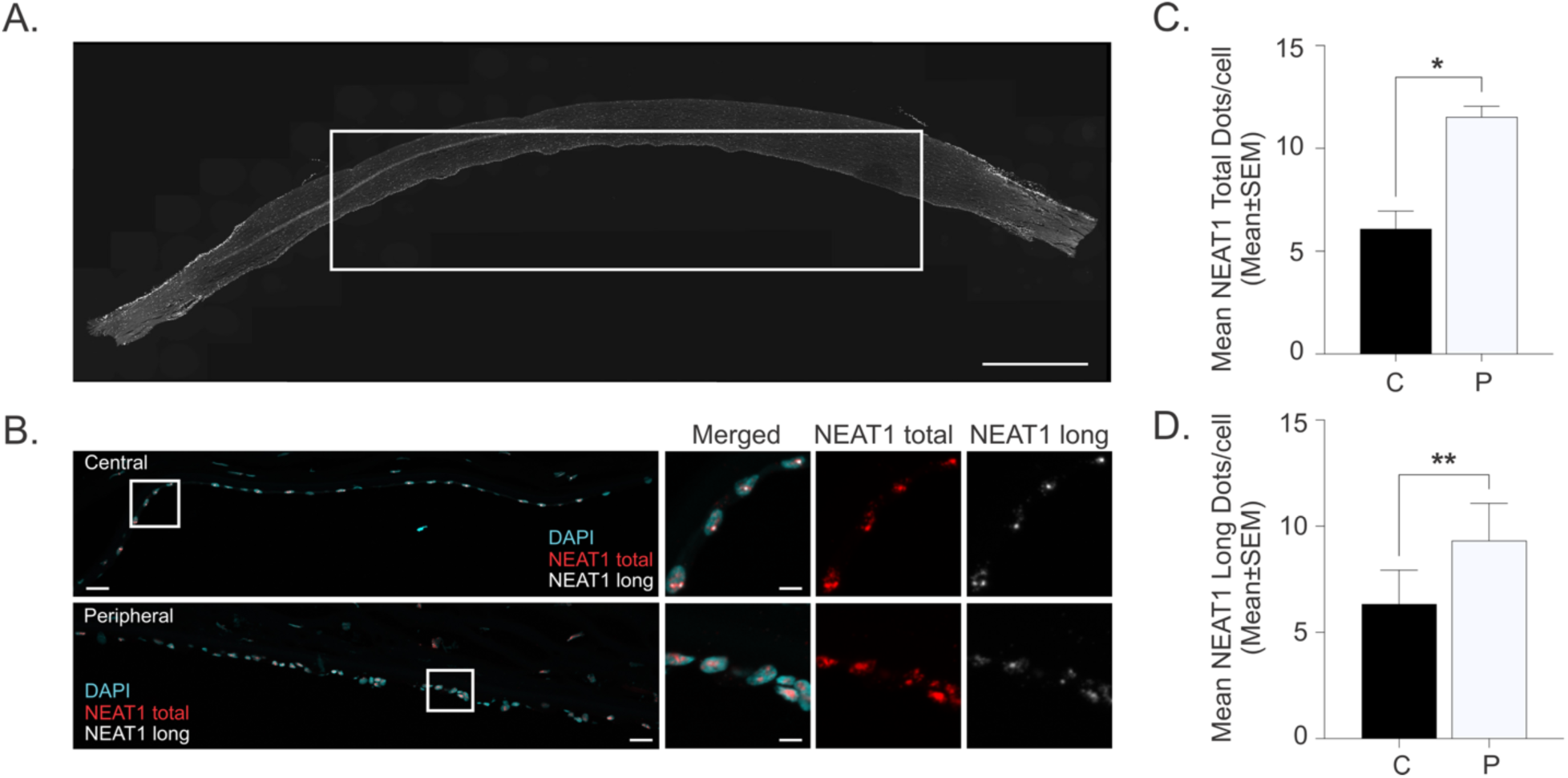
*NEAT1* transcript level is increased in the peripheral corneal endothelium (CE) compared to the central CE. **A)** Representative cross section of a normal cornea (white – DAPI). White box indicates the approximated measured 8-mm central region of the CE layer. (scale bar = 2 mm). **B)** Representative *NEAT1* staining for the central and peripheral spatial domains (scale bar = 50 µm) and magnified images (scale bar = 20 µm) for RNA in situ hybridization staining for the total (short) isoform (red), long isoform (white) and DAPI (cyan). Quantification of fluorescence signal as dots per cell for (**C)** NEAT1 total (short) and **D)** NEAT1 long. *p-value < 0.05; **p-value < 0.01 by two-tailed student *t*-test.

### Elevated NEAT1 levels in normal CECs mitigate oxidative stress mediated-cell death

We assessed endogenous *NEAT1* levels in immortalized cells derived from FECD patients or normal cadaveric donors. *NEAT1* expression was markedly decreased in all 3 FECD CEC lines compared to our normal CECs (Figure 6A). To investigate the response to oxidative stress induced cell death, normal (HCEC-SVN4-68F) and FECD (FECD-SVF7-74F) CECs were stressed with 3.8 mM H_2_O_2_. While normal CECs remained largely unaffected, FECD CECs showed 53.6% decrease in cell viability, reflecting increased sensitivity to H_2_O_2_-mediated oxidative stress in FECD (Figure 6B,C).

**Figure 6.**
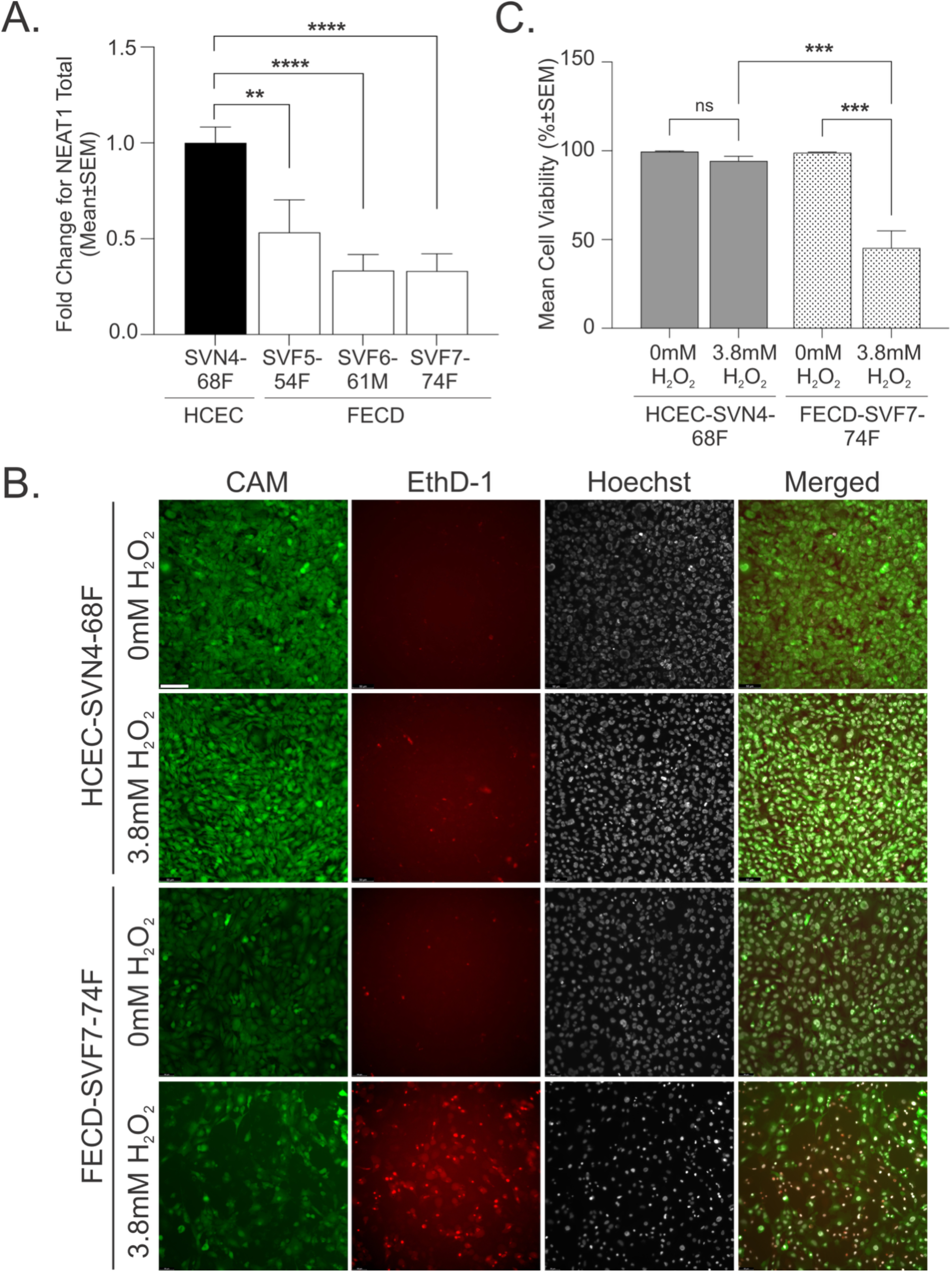
FECD CECs is more susceptible to H_2_O_2_-induced cell death. **A)** Endogenous *NEAT1* levels in normal and FECD immortalized CECs. **p-value < 0.01, ***p-value < 0.001, ****p-value < 0.0001 by one-way ANOVA and Tukey post hoc test. Normal (HCEC-SVN4-68F) and FECD (FECD-SVF7-74F) CECs was treated with 3.8 mM H_2_O_2_. **B)** Representative images of live (green), dead (red) and nuclei (white) staining (scale bar = 50 µm) and **C)** quantification of cell viability are shown. ***p-value < 0.001, ns – not significant by two-way ANOVA and Tukey post hoc test.

### Knockdown of endogenous NEAT1 exacerbates H_2_O_2_-mediated oxidative stress in normal and FECD CECs

To investigate the role of *NEAT1* in regulating the oxidative stress response in CECs, we utilized a NEAT1-targeting shRNA to knockdown *NEAT1* levels in normal and FECD CECs (Figures 7A, S2A). Decrease in *NEAT1* levels, sensitized normal CECs to oxidative stress (Figure 7B). While minimal cell death was observed in normal control CECs (HCEC-SVN4-68F shCTRL), partial *NEAT1* knockdown in normal CECs (HCEC-SVN4-68F shNEAT1) exhibited a 61.6% decreased in cell viability following H_2_O_2_-induced oxidative stress (Figure 7C). Similarly, knockdown of *NEAT1* in FECD CECs, further increased oxidative stress-induced cytotoxicity in this already susceptible cell line (Figure S2B,C). These findings suggest decreased levels of *NEAT1* sensitizes CECs to oxidative stress.

**Figure 7.**
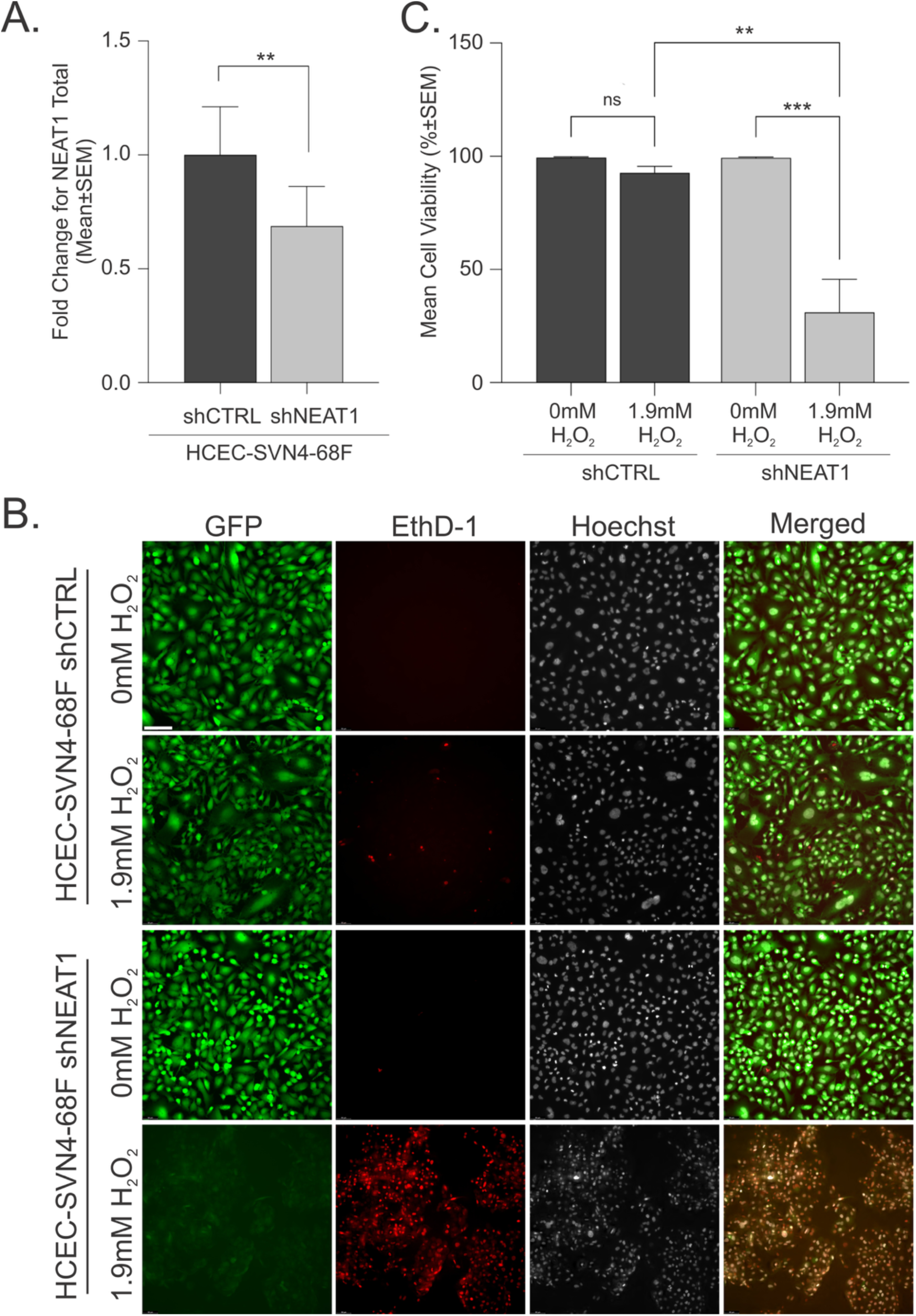
Knockdown of *NEAT1* exacerbates oxidative-stress mediated cell death in normal CECs. **A)** Short hairpin RNA (shRNA) knockdown of *NEAT1* in normal CECs. **p-value < 0.01 by two-tailed student *t*-test. HCEC-SVN4-68F shCTRL and HCEC-SVN4-68F shNEAT1 CECs was treated with 1.9 mM H_2_O_2_. **B)** Representative images of GFP reporter (green), dead (red) and nuclei (white) staining (scale bar = 50 µm) and **C)** quantification of cell viability are shown. **p-value < 0.01, ***p-value < 0.001, ns – not significant by two-way ANOVA and Tukey post hoc test. shCTRL = short hairpin control. sh*NEAT1* = short hairpin *NEAT1*.

### NEAT1 overexpression protects FECD CECs against oxidative stress-induced cell damage

Decreased *NEAT1* levels and the observed increase in cell death following oxidative stress in FECD CECs (Figure 6), prompted us to investigate if we can attenuate H_2_O_2_-induced cell death in FECD CECs. *NEAT1* was overexpressed in normal (HCEC-SVN4-68F) and FECD (FECD-SVF7-74F) CECs (Figures 8A and S3A). In FECD, 66.6% cell death was observed in empty vector control cells (FECD-SVF7-74F EV) following H_2_O_2_-mediated oxidative stress. Remarkably, *NEAT1* overexpression in FECD CECs (FECD-SVF7-74F NEAT1) completely protected the cells from H_2_O_2_-induced cell death (Figure 8B,C). As normal CECs exhibited less sensitivity to H_2_O_2_-induced cell death, *NEAT1*-mediated protection was minimal (Figure S3B,C). These findings demonstrate lncRNA *NEAT1* plays a critical role in protecting CECs from oxidative stress-induced cell damage.

**Figure 8.**
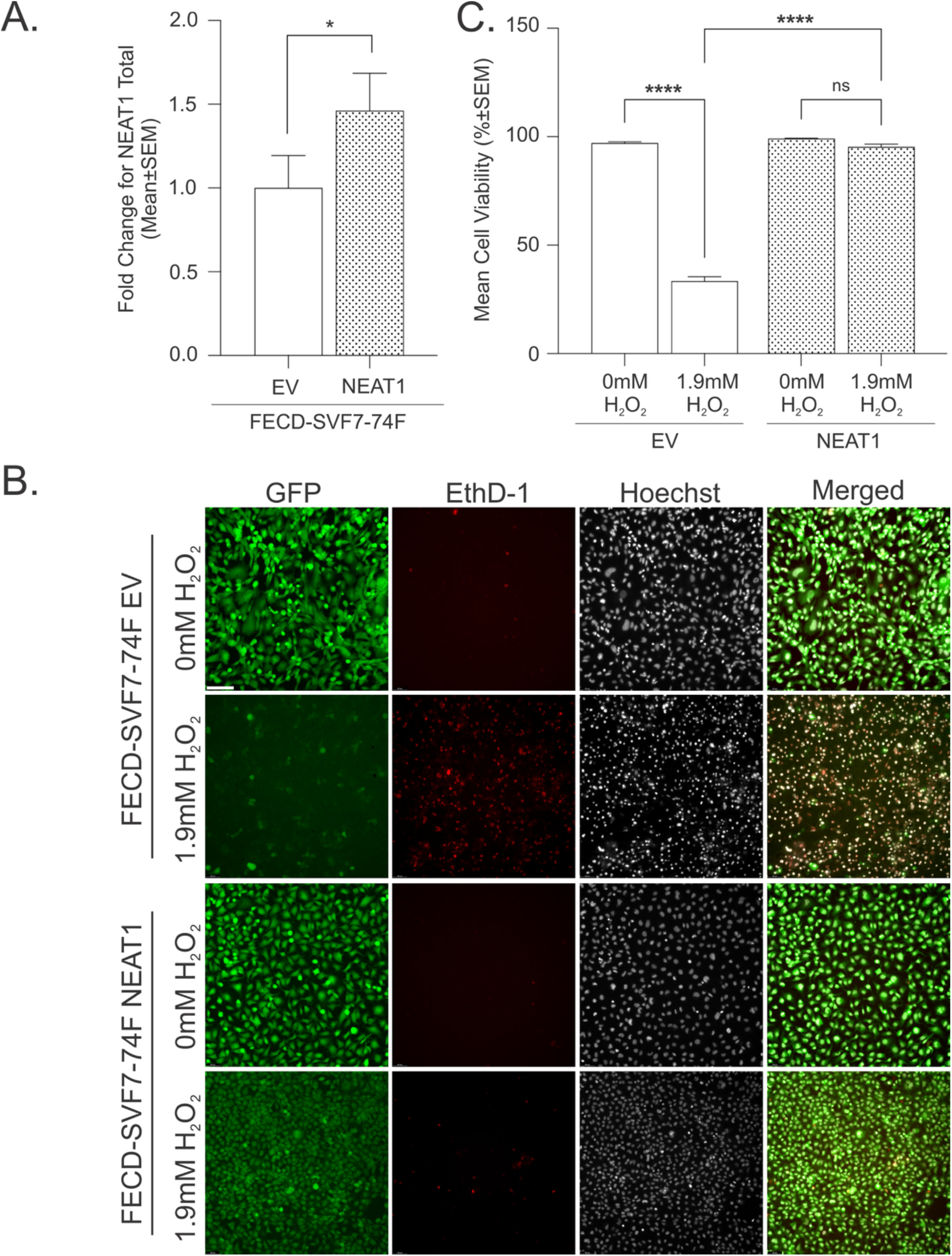
NEAT1 overexpression attenuates oxidative-stress mediated cell death in FECD CECs. **A)** Overexpression of exogenous *NEAT1* in FECD CECs. *p-value < 0.05 by two-tailed student *t*-test. FECD-SVF7-74F EV and FECD-SVF7-74F *NEAT1* CECs was treated with 1.9 mM H_2_O_2_. **B)** Representative images of GFP reporter (green), dead (red) and nuclei (white) staining (scale bar = 50 µm) and **C)** quantification of cell viability are shown. ****p-value < 0.0001, ns – not significant by two-way ANOVA and Tukey post hoc test. EV = empty vector.

## Discussion

The corneal endothelium (CE) is particularly prone to oxidative stress, due to its prolonged exposure to ultraviolet (UV) light, high oxygen and metabolic demand and non-proliferative nature of corneal endothelial cells (CECs). The pathological phenotype observed in FECD begins with the progressive loss of CECs and formation of guttae that is predominately localized to the center of the cornea. As disease progresses, loss of CECs expands outwards from the center to the periphery of the cornea. The mechanisms underlying this spatial pattern of CEC loss, including factors of central versus peripheral susceptibility remains poorly understood and highlights an important clinical gap in our knowledge. Excessive and prolonged exposure to UV radiation in the cornea, in the case of UV-A can cause cellular damage to macromolecules and contribute to increase oxidative stress [20,21]. UV-A increases ROS levels even at low fluences, suggesting UV is especially damaging to cells deficient in antioxidant defense as in FECD [24]. Antioxidant proteins including peroxiredoxins (Prx) are reported to be significantly lower in FECD CECs compared to normal controls [25]. In FECD, CECs are more susceptible to oxidative DNA damage and oxidative stress induced apoptosis than normal CECs, and this likely contributes to the predominately central cell loss and guttae formation observed clinically [24,26,27].

In this study, we show for the first time a key observation that the central CE is more susceptible to H_2_O_2_-mediated cell death compared to the periphery. This increased susceptibility to oxidative stress in the central CE likely contributes to the central pathology observed in FECD. To decipher the spatial difference, we performed RNA sequencing on the central and peripheral CE in normal and FECD cadaveric specimens and identified *NEAT1*, an antioxidant gene, upregulated in the periphery compared to the center of the CE. We further found that *NEAT1* expression was markedly reduced in FECD surgical specimens and CECs, correlating with decreased cell survival following H_2_O_2_-induced oxidative stress relative to normal healthy CECs. Moreover, modulation of *NEAT1* expression directly influenced the cellular response to oxidative injury. Knockdown of *NEAT1* diminished cell viability in normal CECs. More strikingly, increasing *NEAT1* levels in FECD cells fully rescued CECs from H_2_O_2_-induced oxidative stress and cell death, highlighting the importance of *NEAT1*’s role as a cellular protective factor against oxidative damage.

Paraspeckles are nuclear bodies enriched in RNA binding proteins arranged as clusters of foci found in close proximity to nuclear speckles in mammalian cells [31]. Long non-coding RNAs has been found to localize to nuclear bodies, among these is the Nuclear Paraspeckle Assembly Transcript 1 (*NEAT1*) [53]. Despite research attempting to delineate the function of *NEAT1*, its biological role remains to be fully elucidated. In *NEAT1* knockout mouse embryonic fibroblasts, cells were reported to be more sensitive to proteasome inhibitor mediated cell death, suggesting a pro survival role during stress conditions [53]. In murine myeloma cells targeting *NEAT1* downregulates genes involved in DNA damage repair processes [34]. During oxidative stress conditions, *NEAT1* increased viability and decreased apoptosis while exerting a protective effect in blood endothelial cells [40], whereas *NEAT1* was shown to provide a neuroprotective effect against oxidative stress in Parkinson and Huntington’s disease [39,54].

More recently, NEAT1 has been reported to be significantly decreased in FECD CECs [19]. Wang *et al*. [19] reported in a non-genetic FECD animal model and in cell lines, that *NEAT1* knockdown or overexpression exacerbated or attenuated cell death induced by oxidative stress, respectively [19]. However, we did not detect a statistically significant difference in *NEAT1* expression in our RNA sequencing dataset between normal and FECD CE. This discrepancy may reflect the inherent biological heterogeneity of human specimens (our study included all male cadaveric donors), difference in disease severity, variation in sample processing and/or the limited sample size, which may reduce our ability to detect subtle expression changes. However, we consistently detected reduced *NEAT1* levels in both surgical specimens and cell lines derived from patients with a confirmed clinical diagnosis of FECD, supporting previous observations of *NEAT1* dysregulation in FECD. Importantly, we demonstrated for the first time a spatial difference across normal CE with higher *NEAT1* levels in the peripheral region compared to the center. In FECD CE, although a similar trend was observed in the periphery compared to the center this difference did not reach statistical significance, possibly due to our small sample size (N=3). Whether *NEAT1* levels differ within the central and peripheral region with more advanced FECD disease requires further investigation. In line with reports, the knockdown of *NEAT1* increased normal CECs susceptibility to H_2_O_2_-induced oxidative stress. Most notably, the over expression of *NEAT1* in FECD CECs protected the cells from oxidative stress induced cell death; attenuating cell death to levels of the control. Our data along with Wang *et al.* [19] further implicates *NEAT1* as a critical antioxidant in the CE.

The CE has been previously viewed as a homogeneous cell population; however, recent single-cell RNA sequencing analyses identified distinct CEC subsets with unique genetic signatures [19]. The knowledge that the CE is a heterogenous population of cells, together with the identification of putative progenitor-like cells enriched within the inner transition zone at the posterior limbus [18]; raises the intriguing question if this progenitor cell population contributes to the elevated *NEAT1* expression observed in the corneal periphery. In DSO, peripheral CECs migrate centrally to regenerate the central endothelium and restore a functional monolayer. It remains to be determined how *NEAT1* expression changes during this process. This prompts the question of whether migrating peripheral cells retain elevated *NEAT1* expression and how *NEAT1* dynamics evolve as the monolayer re-establishes over time.

FECD is a complex genetic disease with approximately 75% of FECD patients harboring a trinucleotide CTG repeat expansion within the intron region of transcription factor 4 (*TCF4*) [55–57] . Several reports suggest TCF4 and its intronic trinucleotide repeat expansion can modulate the oxidative stress response in FECD [55,58,59]. Saha *et al*. [58] demonstrated FECD surgical samples with unknown TCF4 status and FECD cells harboring the intronic trinucleotide repeat expansion were more susceptible to UVA-mediated ferroptosis as a mechanism of oxidative cell death in FECD [58]. More recently, proteomics analyses on immortalized FECD cells with TCF4 repeat expansion and TCF4-knockout FECD cells revealed significant enrichment in ECM-associated pathways, oxidative stress responses and cellular motility [59]. As CTG repeat expansion status was unknown in our study, whether *NEAT1* levels vary with TCF4 CTG repeat expansion status in FECD and renders CECs more susceptible to oxidative damage remains to be determined.

To our knowledge, this study provides the first evidence that the central CE is intrinsically more vulnerable to oxidative stress, a susceptibility that may be driven in part by reduced *NEAT1* expression. These findings uncover a potential contributor to the characteristic central-to-peripheral progression of FECD. Future studies exploring pharmacological strategies to enhance *NEAT1* expression may open new therapeutic avenues for the treatment of FECD.

## CRediT authorship contribution statement

**Narisa Dhupar:** Formal analysis, Investigation, Validation, Writing – original draft. **Judy Yan:** Investigation, Methodology, Validation, Writing – original draft. **Ness Little:** Investigation. **Stephan Ong Tone:** Conceptualization, Supervision, Funding, Methodology, Project Administration, Writing – review & editing.

## Data availability

Data available upon request.

## Acknowledgements

We would like to thank Dr. Ula Jurkunas at the Schepens Eye Research Institute in Boston, Massachusetts for graciously providing the cell lines and Dr. Chao Wang for her expertise and comments on the manuscript.

---

**Figure S1: Specular micrographs for FECD specimens.** Representative images confirming FECD diagnosis, showing the corneal endothelium (white) and characteristic guttae (dark spots) in FECD_1 and FECD_3 specimens. A clinical diagnosis of FECD was confirmed through donor medical history for FECD_2.

**Figure S2. Decrease in NEAT1 expression enhances hydrogen peroxide-mediated cell death in FECD CECs. A)** *NEAT1* levels following shRNA knockdown in FECD CECs. ***p < 0.001 by two-tailed student *t*-test. FECD-SVF7-74F shCTRL and FECD-SVF7-74F shNEAT1 CECs was treated with 1.9 mM H_2_O_2_. **B)** Representative images of GFP reporter (green), dead (red) and nuclei (white) staining (scale bar = 50 µm) and **C)** quantification of cell viability are shown. *p-value < 0.05, ****p-value < 0.0001 by two-way ANOVA and Tukey post hoc test. shCTRL = short hairpin control. sh*NEAT1* = short hairpin *NEAT1*.

**Figure S3. NEAT1 overexpression attenuates hydrogen peroxide-mediated cell death in normal CECs. A)** Overexpression of exogenous *NEAT1* in normal CECs. *p-value < 0.05 by two-tailed student *t*-test. HCEC-SVN4-68F EV and HCEC-SVN4-68F *NEAT1* CECs was treated with 1.9 mM H_2_O_2_. **B)** Representative images of GFP reporter (green), dead (red) and nuclei (white) staining (scale bar = 50 µm) and **C)** quantification of cell viability are shown. ***p < 0.001, ****p-value < 0.0001, ns – not significant by two-way ANOVA and Tukey post hoc test. EV = empty vector.

