## Supplementary figures and images for "Differential expression of *NEAT1* in the corneal endothelium increases susceptibility to oxidative stress in Fuchs Endothelial Corneal Dystrophy"

### Supplementary Figures

#### Figure S1

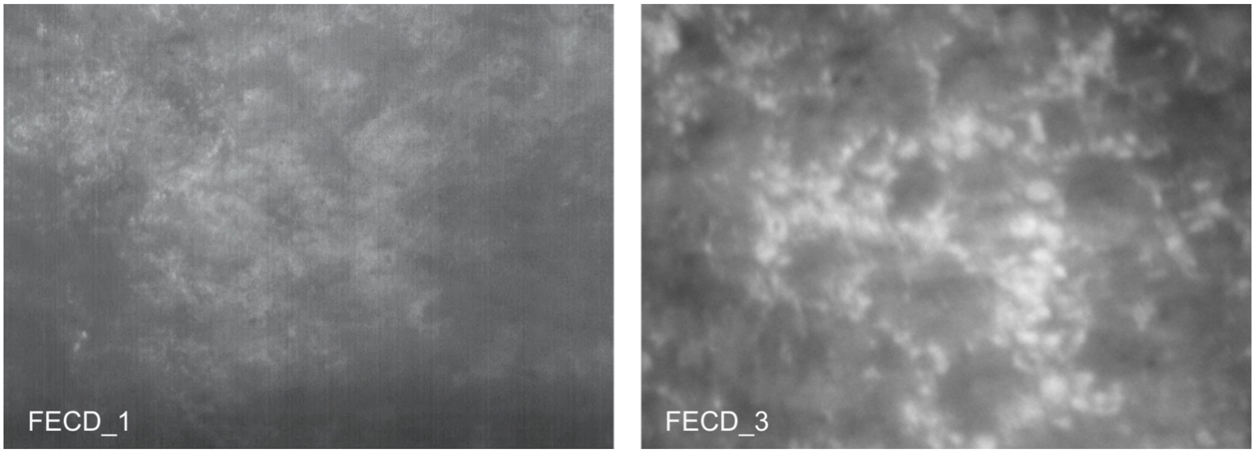

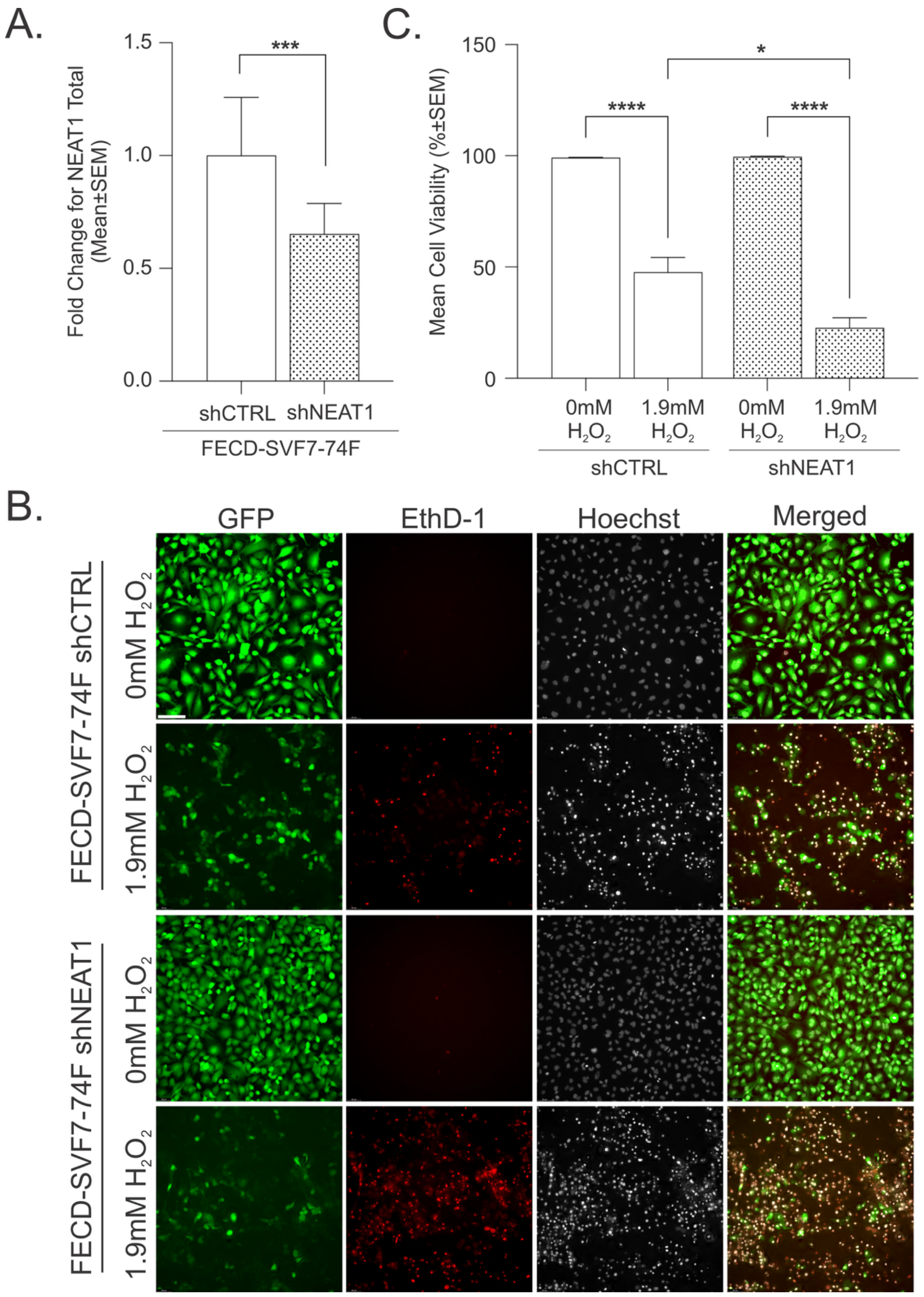

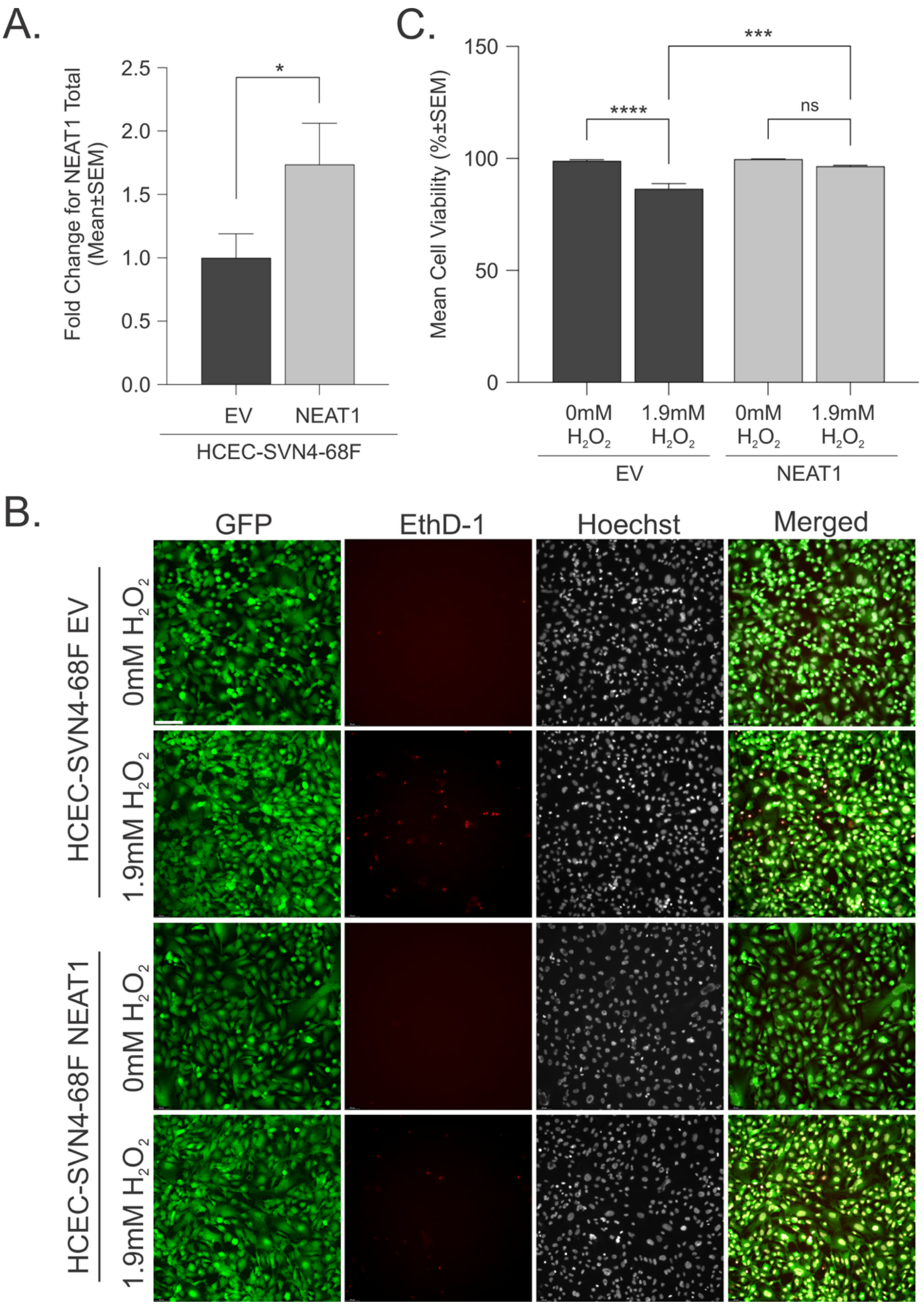
