## Supplementary Tables for "Differential expression of *NEAT1* in the corneal endothelium increases susceptibility to oxidative stress in Fuchs Endothelial Corneal Dystrophy"

**Supplementary Table 1.** Patient and Donor Specimen Characteristics

| **CategoryΔ** | **Tissue ID⊥** | **Laterality✱** | **Age (yr)** | **Sex** | **Cause of Death** | **Death to Preservation Time (hrs:mins)** | **RNA Concentration (ng/uL)✚** | **RIN†** | **Experiment** |
|---|---|---|---|---|---|---|---|---|---|
| Normal | N_1 | OS | 74 | Male | COPD, Right Lower Lobe Pneumonia | 8:30 | **C:** 33.4 **P:** 28.6 | **C:** 8.5 **P:** 7.3 | RNA-seq |
| Normal | N_2 | OD | 71 | Female | Anoxia, Pneumonia, CHF | 22:20 | **C:** 59.0 **P:** 37.4 | **C:** 6.9 **P:** 7.3 | RNA-seq |
| Normal | N_3 | OD | 60 | Male | MI, Atherosclerosis | 24:32 | **C:** 32.6 **P:** 40.6 | **C:** 8.4 **P:** 6.2 | RNA-seq |
| Normal | N_4 | OS | 49 | Female | Cardiac Arrest | 18:37 | - | - | $H_2O_2$ Assay |
| Normal | N_5 | OS | 76 | Male | Obstructed Tracheostomy Tube | 17:45 | - | - | $H_2O_2$ Assay |
| Normal | N_6 | OS | 73 | Male | Cardiogenic Shock | 20:32 | - | - | $H_2O_2$ Assay |
| Normal | N_7 | OS | 79 | Female | Cancer | 16:20 | - | - | $H_2O_2$ Assay |
| Normal | N_8 | OD | 76 | Male | PE, CAD | 22:01 | - | - | $H_2O_2$ Assay |
| Normal | N_9 | OS | 76 | Male | PE, CAD | 22:01 | - | - | $H_2O_2$ Assay |
| Normal | N_10 | OS | 74 | Female | Peritonitis | 7:32 | - | - | $H_2O_2$ Assay |
| Normal | N_11 | OD | 51 | Female | Cerebrovascular Accident, Stroke | 25:05 | - | - | $H_2O_2$ Assay |
| Normal | N_12 | OD | 49 | Female | Cardiac Arrest | 18:37 | - | - | RNAscope |
| Normal | N_13 | OD | 76 | Male | Obstructed Tracheostomy Tube | 17:45 | - | - | RNAscope |
| Normal | N_14 | OD | 79 | Female | Cancer | 16:20 | - | - | RNAscope |
| Normal | N_15 | OS | 65 | Female | Metabolic Acidosis | 9:57 | - | - | qRT-PCR |
| Normal | N_16 | OS | 62 | Male | Cardiac Dysrhythmia, Cancer | 23:36 | - | - | qRT-PCR |
| Normal | N_17 | OS | 74 | Female | COPD | 20:24 | - | - | qRT-PCR |
| Normal | N_18 | OS | 68 | Male | Cardiac Arrest | 22:11 | - | - | qRT-PCR |
| Normal | N_19 | OS | 76 | Male | Sepsis | 7:50 | - | - | qRT-PCR |
| Normal | N_20 | OS | 73 | Male | Cancer | 18:50 | - | - | qRT-PCR |
| Normal | N_21 | OS | 54 | Male | Cardiac Arrest, Anoxia | 12:40 | - | - | qRT-PCR |
| Normal | N_22 | OS | 79 | Male | Pneumonia, Respiratory Failure | 9:56 | - | - | qRT-PCR |
| Normal | N_23 | OS | 69 | Male | Cancer | 11:21 | - | - | qRT-PCR |
| FECD | FECD_1 | OD | 75 | Male | CAD | 8:18 | **C:** 18.5 **P:** 18.8 | **C:** 8.0 **P:** 7.0 | RNA-seq |
| FECD | FECD_2 | OD | 69 | Female | Spinal Stenosis, MAID | 12:03 | **C:** 27.4 **P:** 33.4 | **C:** 7.3 **P:** 6.7 | RNA-seq |
| FECD | FECD_3 | OS | 66 | Male | MI | 16:11 | **C:** 22.4 **P:** 25.0 | **C:** 6.8 **P:** 4.9 | RNA-seq |
| FECD | FECD_4 | OS | 55 | Female | - | - | - | - | qRT-PCR |
| FECD | FECD_5 | OD | 82 | Male | - | - | - | - | qRT-PCR |
| FECD | FECD_6 | OS | 70 | Male | - | - | - | - | qRT-PCR |
| FECD | FECD_7 | OD | 55 | Male | - | - | - | - | qRT-PCR |
| FECD | FECD_8 | OS | 77 | Female | - | - | - | - | qRT-PCR |
| FECD | FECD_9 | OS | 75 | Male | - | - | - | - | qRT-PCR |

Δ FECD_4 to FECD_9 surgical specimens with confirmed clinical FECD diagnosis

⊥N=Normal; FECD=Fuchs Endothelial Corneal Dystrophy

✱OS=Left Eye; OD=Right Eye

✚RNA concentrations measured by NanoDrop 2000; C=Central Endothelium; P=Peripheral Endothelium.

†RIN=RNA Integrity Number

COPD= Chronic Obstructive Pulmonary Disease, PE=Pulmonary Embolism, CAD=Coronary Artery Disease, MAID=Medical Assistance in Dying, MI=Myocardial Infraction, CHF=Congestive Heart Failure

**Supplementary Table 2.** qRT-PCR Primer Sequences

| **Primers** | **Sequence 5'-3'** |
| --- | --- |
| NEAT1 long | **Forward:** GGCCAGAGCTTTGTTGCTTC<br>**Reverse:** GGTGCGGGCACTTACTTACT |
| NEAT1 short | **Forward:** TGGCTAGCTCAGGGCTTCAG<br>**Reverse:** TCTCCTTGCCAAGCTTCCTTC |
| β-Actin | **Forward:** ACCGAGCGCGGCTACAG<br>**Reverse:** CTTAATGTCACGCACGATTTCC |

**Supplementary Table 3.** Full list of upregulated and downregulated differentially expressed genes in a healthy corneal endothelium between the central and the peripheral spatial domains.

| **Upregulated in Periphery** | | | **Downregulated in Periphery** | | |
|---|---|---|---|---|---|
| **Gene Symbol** | **log 2FC (P/C) †** | **ID** | **Gene Symbol** | **log 2FC (P/C) †** | **ID** |
| RP11-707G18.1 | 0.65075882 | ENSG00000278743 | EXD2 | -2.1267956 | ENSG00000081177 |
| GNG11 | 1.49057859 | ENSG00000127920 | ELP6 | -0.8920773 | ENSG00000163832 |
| AC093838.4 | 0.60640956 | ENSG00000152117 | CCL28 | -0.7965152 | ENSG00000151882 |
| FLNA | 0.88624577 | ENSG00000196924 | EDN3 | -1.5441259 | ENSG00000124205 |
| ZNF503 | 2.006034736 | ENSG00000165655 | TBC1D4 | -0.6268893 | ENSG00000136111 |
| GABBR1 | 0.97733336 | ENSG00000204681 | TMEM45B | -0.7050558 | ENSG00000151715 |
| DHX36 | 1.16690431 | ENSG00000174953 | CFH | -0.9680763 | ENSG00000000971 |
| RN7SL653P | 0.88231372 | ENSG00000239794 | FGF9 | -1.3867984 | ENSG00000102678 |
| STAB1 | 1.51119668 | ENSG00000010327 | ING4 | -0.7641795 | ENSG00000111653 |
| RP11-186B7.4 | 0.87418212 | ENSG00000264772 | CFHR1 | -0.7360377 | ENSG00000244414 |
| RHBDF1 | 0.69940174 | ENSG00000007384 | CA3 | -0.8216713 | ENSG00000164879 |
| CTD-3252C9.4 | 1.3449358 | ENSG00000267519 | PAPPA | -0.6683357 | ENSG00000182752 |
| RASA3 | 1.13722841 | ENSG00000185989 | EIF2B5 | -1.0603036 | ENSG00000145191 |
| C5orf45 | 0.72863297 | ENSG00000161010 | RP11-205K6.3 | -0.9262743 | ENSG00000279137 |
| RP11-181G12.2 | 1.35556712 | ENSG00000182873 | TM2D1 | -0.6056064 | ENSG00000162604 |
| COL3A1 | 1.79375571 | ENSG00000168542 | OR51E2 | -0.6771846 | ENSG00000167332 |
| RP11-73M18.6 | 0.88062925 | ENSG00000270108 | ARHGDIB | -0.9122549 | ENSG00000111348 |
| NEAT1 | 1.75864001 | ENSG00000245532 | ABHD14A-ACY1 | -1.2185768 | ENSG00000114786 |
| MIR4639 | 0.89475451 | ENSG00000263712 | RPE | -0.6641437 | ENSG00000197713 |
| PABPC1L | 0.60311255 | ENSG00000101104 | E2F5 | -0.6811843 | ENSG00000133740 |
| AF011889.5 | 0.65958364 | ENSG00000241489 | RP11-334C17.6 | -0.8721455 | ENSG00000275479 |

| | | | | | |
|---|---|---|---|---|---|
| EIF2AK3 | 0.59467446 | ENSG00000172071 | PTRH2 | -0.6302022 | ENSG00000141378 |
| PHYKPL | 1.26404767 | ENSG00000175309 | RPS3 | -0.6259779 | ENSG00000149273 |
| NAA25 | 0.68839123 | ENSG00000111300 | KHDRBS3 | -0.7480684 | ENSG00000131773 |
| MT1E | 1.942834 | ENSG00000169715 | CSNK1G3 | -1.0347231 | ENSG00000151292 |
| FEZ1 | 1.36088625 | ENSG00000149557 | SLC37A4 | -0.9899587 | ENSG00000137700 |
| SLC25A27 | 0.6089601 | ENSG00000153291 | TEN1 | -1.2435685 | ENSG00000257949 |
| AC092910.1 | 1.19153302 | ENSG00000276607 | EARS2 | -0.6474052 | ENSG00000103356 |
| snoU13 | 0.79547585 | ENSG00000238311 | CDKN2AIPNL | -0.6504868 | ENSG00000237190 |
| HLA-C | 0.67355147 | ENSG00000204525 | RP11-205K6.1 | -1.0535883 | ENSG00000233721 |
| TAC1 | 3.48010072 | ENSG00000006128 | TPRKB | -1.1176368 | ENSG00000144034 |
| NPDC1 | 1.10420726 | ENSG00000107281 | MPP1 | -0.7626223 | ENSG00000130830 |
| BABAM1 | 0.70485662 | ENSG00000105393 | NUAK1 | -0.7590057 | ENSG00000074590 |
| SEMA3F | 1.33961533 | ENSG00000001617 | PDHA1 | -0.6265166 | ENSG00000131828 |
| AASS | 0.66180827 | ENSG00000008311 | SULT1A1 | -1.1589335 | ENSG00000196502 |
| AP001205.1 | 1.56800041 | ENSG00000264991 | TMEM55B | -0.7194756 | ENSG00000165782 |
| YBX3 | 1.73832608 | ENSG00000060138 | POLR2B | -0.6072099 | ENSG00000047315 |
| TGIF1 | 0.75801147 | ENSG00000177426 | ITGAE | -0.6536511 | ENSG00000083457 |
| EPOR | 1.0271039 | ENSG00000187266 | AMPD3 | -0.9979499 | ENSG00000133805 |
| RP3-368A4.6 | 0.94137918 | ENSG00000271533 | ATP6V0A4 | -1.082069 | ENSG00000105929 |
| SIRPA | 0.82002893 | ENSG00000198053 | HEXIM2 | -0.908636 | ENSG00000168517 |
| RSRP1 | 1.49762786 | ENSG00000117616 | NPC2 | -0.7398032 | ENSG00000119655 |
| PCSK1N | 1.59734797 | ENSG00000102109 | NEDD4L | -0.7180011 | ENSG00000049759 |
| SMG1P3 | 0.92354591 | ENSG00000180747 | SNHG21 | -0.6367333 | ENSG00000250988 |
| SERPING1 | 0.66359262 | ENSG00000149131 | KISS1 | -1.1658965 | ENSG00000170498 |
| ACHE | 1.67420428 | ENSG00000087085 | FAM193A | -0.9569667 | ENSG00000125386 |
| COL5A1 | 1.38929019 | ENSG00000130635 | FRMPD1 | -0.7643779 | ENSG00000070601 |

| | | | | | |
|---|---|---|---|---|---|
| HAUS5 | 0.97232708 | ENSG00000249115 | ENO1-AS1 | -0.6705544 | ENSG00000230679 |
| COL6A2 | 3.01747399 | ENSG00000142173 | MRPL22 | -1.4862281 | ENSG00000082515 |
| MIR7152 | 2.03867682 | ENSG00000274824 | PFDN6 | -0.6463367 | ENSG00000204220 |
| RP11-373D23.2 | 0.74829797 | ENSG00000270640 | CDK5 | -0.9189067 | ENSG00000164885 |
| ZNF791 | 0.59092247 | ENSG00000173875 | FAM89A | -0.6470832 | ENSG00000182118 |
| IL13RA2 | 1.21133575 | ENSG00000123496 | LRRC23 | -0.8388263 | ENSG00000010626 |
| RN7SL600P | 0.97070721 | ENSG00000274963 | EBP | -0.6761202 | ENSG00000147155 |
| COL1A1 | 2.13840772 | ENSG00000108821 | CYP51A1 | -0.6221467 | ENSG00000001630 |
| ANKRD36 | 0.67952985 | ENSG00000135976 | ISM2 | -1.2314412 | ENSG00000100593 |
| ARHGEF1 | 0.80308091 | ENSG00000076928 | MPP5 | -0.8697817 | ENSG00000072415 |
| CCNL2 | 0.59256858 | ENSG00000221978 | PPP1R3G | -0.6297358 | ENSG00000219607 |
| DDIT4L | 1.31406906 | ENSG00000145358 | ZSCAN16-AS1 | -0.8703805 | ENSG00000269293 |
| FAT4 | 1.15160524 | ENSG00000196159 | NECAB1 | -0.8993474 | ENSG00000123119 |
| USP32P2 | 0.95269225 | ENSG00000233327 | KAZN | -0.6843649 | ENSG00000189337 |
| USP40 | 0.76640486 | ENSG00000085982 | OSBPL1A | -0.6502682 | ENSG00000141447 |
| GYPC | 0.93185229 | ENSG00000136732 | AC007405.6 | -0.7080494 | ENSG00000239467 |
| AC091053.2 | 0.84952994 | ENSG00000281902 | GGCT | -0.8853956 | ENSG00000006625 |
| ENOSF1 | 0.66018181 | ENSG00000132199 | ZSWIM1 | -0.9377963 | ENSG00000168612 |
| FTX | 0.81373356 | ENSG00000230590 | RORC | -0.7523011 | ENSG00000143365 |
| NES | 1.65662005 | ENSG00000132688 | RPP30 | -1.0396268 | ENSG00000148688 |
| FAM187A | 1.16087666 | ENSG00000214447 | ALG8 | -0.6394031 | ENSG00000159063 |
| SOD3 | 1.40350972 | ENSG00000109610 | COX20 | -0.6145682 | ENSG00000203667 |
| LAYN | 0.7796243 | ENSG00000204381 | MOCS2 | -0.6579496 | ENSG00000164172 |
| EHBP1L1 | 1.00516216 | ENSG00000173442 | S100A4 | -0.6773826 | ENSG00000196154 |
| RP11-727A23.4 | 0.82360601 | ENSG00000254676 | DEPTOR | -0.8090494 | ENSG00000155792 |
| BEX1 | 1.31108807 | ENSG00000133169 | RP11-140K17.3 | -1.0686928 | ENSG00000272288 |

| | | | | | |
|---|---|---|---|---|---|
| PLA2G4B | 0.80760004 | ENSG00000243708 | METTL23 | -0.6805516 | ENSG00000181038 |
| MALAT1 | 0.7957549 | ENSG00000251562 | ERBB3 | -0.6493142 | ENSG00000065361 |
| SMG1P4 | 0.82386923 | ENSG00000185710 | PDHB | -0.6627236 | ENSG00000168291 |
| ADAMTSL4-AS1 | 0.74218381 | ENSG00000203804 | RP5-834N19.1 | -0.8327266 | ENSG00000232650 |
| ABCA7 | 0.60869437 | ENSG00000064687 | C9orf116 | -0.6464102 | ENSG00000160345 |
| MIR6777 | 0.76555089 | ENSG00000274111 | NAT1 | -0.9206882 | ENSG00000171428 |
| ITPR3 | 1.97762421 | ENSG00000096433 | CTD-3214H19.4 | -1.053838 | ENSG00000268400 |
| FZD1 | 0.7578455 | ENSG00000157240 | RP11-321G12.1 | -0.6620778 | ENSG00000259459 |
| ZNF638-IT1 | 0.65510441 | ENSG00000281195 | | | |
| FSTL3 | 0.7247764 | ENSG00000070404 | | | |
| AHSA2 | 0.64839149 | ENSG00000173209 | | | |
| DUS1L | 0.78514108 | ENSG00000169718 | | | |

† Log2 Fold Change (Peripheral/Central).

**Supplementary Table 4.** Full list of upregulated and downregulated differentially expressed genes in a FECD corneal endothelium between the central and the peripheral spatial domains

| Upregulated in Periphery | | | Downregulated in Periphery | | |
|---|---|---|---|---|---|
| Gene Symbol | $log_2FC$ (P/C) † | ID | Gene Symbol | $log_2FC$ (P/C) † | ID |
| ELN | 2.08108576 | ENSG00000049540 | EXOSC8 | -0.9663734 | ENSG00000120699 |
| KIF1A | 2.42960025 | ENSG00000130294 | TXNRD3 | -1.6523686 | ENSG00000197763 |
| MIR6809 | 1.63823689 | ENSG00000275458 | RASSF6 | -1.5693554 | ENSG00000169435 |
| Metazoa_SRP | 0.87936923 | ENSG00000278771 | MEGF9 | -0.8523942 | ENSG00000106780 |
| KIAA1683 | 1.64110132 | ENSG00000130518 | PTS | -0.6009438 | ENSG00000150787 |
| Metazoa_SRP | 1.01203565 | ENSG00000274012 | LIG4 | -1.0168507 | ENSG00000174405 |
| RP5-940J5.9 | 2.47645233 | ENSG00000269968 | ITGA1 | -0.9157934 | ENSG00000213949 |
| SRRM3 | 1.62587047 | ENSG00000177679 | MAGED4B | -1.0120243 | ENSG00000187243 |
| TMEM132C | 2.14686485 | ENSG00000181234 | TFRC | -0.8199718 | ENSG00000072274 |
| SFRP4 | 1.84863447 | ENSG00000106483 | TENM3 | -1.1533319 | ENSG00000218336 |
| ADAMTSL3 | 3.06468033 | ENSG00000156218 | NECAB1 | -0.6283738 | ENSG00000123119 |
| RSPO4 | 3.22295337 | ENSG00000101282 | GNG12 | -0.8050425 | ENSG00000172380 |
| SRPX | 1.72721548 | ENSG00000101955 | USP10 | -0.623766 | ENSG00000103194 |
| PLA2G2A | 3.88551732 | ENSG00000188257 | DUSP14 | -0.6605848 | ENSG00000276023 |
| SEPT4-AS1 | 1.32356535 | ENSG00000264672 | TIMP2 | -0.6139097 | ENSG00000035862 |
| PTN | 3.49014335 | ENSG00000105894 | PAPPA | -0.961848 | ENSG00000182752 |
| HMGN2P15 | 1.36601703 | ENSG00000214578 | TMPRSS3 | -1.1019157 | ENSG00000160183 |
| ACHE | 1.62074722 | ENSG00000087085 | GPNMB | -0.761496 | ENSG00000136235 |
| SCARNA6 | 1.66504577 | ENSG00000251791 | SCNM1 | -0.8827325 | ENSG00000163156 |
| MAPKBP1 | 0.61578129 | ENSG00000137802 | PTPRK | -0.7042892 | ENSG00000152894 |
| PRX | 0.71688564 | ENSG00000105227 | RHOBTB2 | -0.616087 | ENSG00000008853 |

| | | | | | |
|---|---|---|---|---|---|
| SNORD17 | 1.35434128 | ENSG00000212232 | DMTN | -1.0573139 | ENSG00000158856 |
| AC002044.1 | 0.85417545 | ENSG00000279591 | FKTN | -1.2879714 | ENSG00000106692 |
| RBP4 | 3.38511423 | ENSG00000138207 | PRKG1 | -0.7913648 | ENSG00000185532 |
| GUCY1A3 | 2.87741843 | ENSG00000164116 | ZFYVE27 | -0.6834598 | ENSG00000155256 |
| MLXIPL | 1.55484388 | ENSG00000009950 | DCK | -0.905247 | ENSG00000156136 |
| RN7SL1 | 0.82860106 | ENSG00000276168 | | | |
| RSPO2 | 3.34173482 | ENSG00000147655 | | | |
| SMOC2 | 3.96701826 | ENSG00000112562 | | | |
| RP11-554D20.2 | 0.7498499 | ENSG00000273792 | | | |
| FGFR3 | 1.92771783 | ENSG00000068078 | | | |
| TYRP1 | 2.35478532 | ENSG00000107165 | | | |
| FBLN1 | 3.39626098 | ENSG00000077942 | | | |
| SDPR | 1.98306186 | ENSG00000168497 | | | |
| RN7SL4P | 1.03675926 | ENSG00000263740 | | | |
| DANT2 | 0.61167948 | ENSG00000235244 | | | |
| ISLR | 2.53864383 | ENSG00000129009 | | | |
| RMST | 2.24196927 | ENSG00000255794 | | | |
| FXYD1 | 2.05179291 | ENSG00000266964 | | | |
| CCDC144A | 0.94524359 | ENSG00000170160 | | | |
| COL16A1 | 3.1402109 | ENSG00000084636 | | | |
| CLMN | 1.33305191 | ENSG00000165959 | | | |
| PTPRF | 0.82740215 | ENSG00000142949 | | | |
| FAM13C | 1.40081935 | ENSG00000148541 | | | |
| RP11-385D13.3 | 1.29574737 | ENSG00000266538 | | | |
| RN7SL2 | 1.08466243 | ENSG00000280102 | | | |
| RP11-588K22.2 | 2.4580181 | ENSG00000260244 | | | |

| | | |
|---|---|---|
| MIR6852 | 1.10056482 | ENSG00000275831 |
| MIR770 | 2.48992217 | ENSG00000211574 |
| AC025279.1 | 1.19392983 | ENSG00000278509 |
| CHI3L1 | 3.32661584 | ENSG00000133048 |
| SLC24A3 | 2.00332229 | ENSG00000185052 |
| LMOD1 | 2.10547609 | ENSG00000163431 |
| SLC35D1 | 0.72527372 | ENSG00000116704 |
| RN7SL674P | 0.74264661 | ENSG00000239899 |
| TIAM1 | 2.01113729 | ENSG00000156299 |
| FAM129A | 1.7915854 | ENSG00000135842 |
| CPSF3L | 0.60029028 | ENSG00000127054 |
| RP11-326C3.7 | 1.15136762 | ENSG00000254910 |
| KCNE4 | 2.63863709 | ENSG00000152049 |
| STAB1 | 2.2394623 | ENSG00000010327 |
| EMP2 | 1.73776272 | ENSG00000213853 |
| C10orf10 | 2.11090549 | ENSG00000165507 |
| ADAMTS10 | 1.54508751 | ENSG00000142303 |
| MIR2278 | 1.08367243 | ENSG00000252153 |
| CTTNBP2 | 1.75083898 | ENSG00000077063 |
| GPR162 | 0.76976478 | ENSG00000250510 |
| XAF1 | 1.06111988 | ENSG00000132530 |
| mascRNA-menRNA | 3.18557065 | ENSG00000275948 |
| SNORD116-1 | 0.92954466 | ENSG00000207063 |
| RP11-54A4.2 | 1.33430518 | ENSG00000237781 |
| KCNAB1 | 1.73120863 | ENSG00000169282 |
| CTD-3035K23.7 | 1.27638279 | ENSG00000274565 |

| | | |
|---|---|---|
| NEFL | 1.63954416 | ENSG00000277586 |
| SNORA73A | 1.84430761 | ENSG00000274266 |
| AEBP1 | 3.03357552 | ENSG00000106624 |
| RP11-228B15.4 | 0.80981941 | ENSG00000225032 |
| RN7SL600P | 0.99252973 | ENSG00000274963 |
| CDH23 | 2.10565187 | ENSG00000107736 |
| RIMS4 | 1.23737167 | ENSG00000101098 |
| TMEFF2 | 1.87309926 | ENSG00000144339 |
| CMKLR1 | 1.30485698 | ENSG00000174600 |
| PDGFRA | 3.1935378 | ENSG00000134853 |
| SNORA54 | 1.64376763 | ENSG00000207008 |
| SCARNA12 | 1.76844098 | ENSG00000238795 |
| GUSBP11 | 1.14503494 | ENSG00000228315 |
| FIBIN | 2.22569736 | ENSG00000176971 |
| LIFR | 2.89486889 | ENSG00000113594 |
| ALDH1A1 | 2.20273744 | ENSG00000165092 |
| SLC41A2 | 0.9497959 | ENSG00000136052 |
| MXRA5 | 3.44460704 | ENSG00000101825 |
| PLP1 | 2.19721295 | ENSG00000123560 |
| GAS7 | 1.90201423 | ENSG00000007237 |
| WDR86 | 1.503426 | ENSG00000187260 |
| SCARNA13 | 1.17434141 | ENSG00000252481 |
| RP11-379F4.1 | 1.09457008 | ENSG00000243675 |
| ANGPTL7 | 4.01249429 | ENSG00000171819 |
| THBS2 | 1.11836509 | ENSG00000186340 |
| PENK | 2.49251516 | ENSG00000181195 |

| | | |
|---|---|---|
| SDR39U1 | 0.91541725 | ENSG00000100445 |
| MAPK15 | 1.37263339 | ENSG00000181085 |
| SLC8A3 | 1.08841551 | ENSG00000100678 |
| MAST2 | 0.93721735 | ENSG00000086015 |
| SYNPO2 | 1.67628252 | ENSG00000172403 |
| PCSK2 | 1.25772115 | ENSG00000125851 |
| CLMP | 2.01292972 | ENSG00000166250 |
| MIR590 | 0.84215892 | ENSG00000207741 |
| SCARNA5 | 1.00820357 | ENSG00000252010 |
| C1R | 3.25121921 | ENSG00000159403 |
| TMEM176B | 2.51204344 | ENSG00000106565 |
| SAMHD1 | 1.55174719 | ENSG00000101347 |
| RP3-449O17.1 | 0.88670828 | ENSG00000244627 |
| HLA-F | 1.41647142 | ENSG00000204642 |
| SCARNA21 | 0.89974149 | ENSG00000252835 |
| DOC2B | 2.64970122 | ENSG00000272636 |
| ENOSF1 | 0.80703262 | ENSG00000132199 |
| SCARA5 | 2.05341841 | ENSG00000168079 |
| CPE | 2.80838519 | ENSG00000109472 |
| PDE1C | 1.47783594 | ENSG00000154678 |
| SLC2A9 | 1.18677613 | ENSG00000109667 |
| TLE2 | 2.39807657 | ENSG00000065717 |
| GUCY1B3 | 1.59524541 | ENSG00000061918 |
| ANKRD36C | 0.78184906 | ENSG00000174501 |
| MEG3 | 2.77677508 | ENSG00000214548 |
| BMPER | 1.85221278 | ENSG00000164619 |

| | | |
|---|---|---|
| RMRP | 1.78365332 | ENSG00000277027 |
| RN7SL648P | 1.4090486 | ENSG00000265727 |
| C1S | 2.0634157 | ENSG00000182326 |
| HSD11B1 | 2.35246377 | ENSG00000117594 |
| TNFRSF21 | 1.78831922 | ENSG00000146072 |
| AC106788.1 | 1.85169898 | ENSG00000276305 |
| AC133555.1 | 1.53625949 | ENSG00000276338 |
| PRRG4 | 1.40214795 | ENSG00000135378 |
| SLIT3 | 1.48870979 | ENSG00000184347 |
| IGFBP6 | 1.69689069 | ENSG00000167779 |
| MIR6775 | 1.58225406 | ENSG00000278598 |
| FAM167B | 0.92889369 | ENSG00000183615 |
| RP11-47G11.2 | 1.11645581 | ENSG00000232334 |
| PMEL | 1.67044844 | ENSG00000185664 |
| DNM1P47 | 0.89947637 | ENSG00000259660 |
| TMEM173 | 1.13871277 | ENSG00000184584 |
| RAMP2 | 1.53640248 | ENSG00000131477 |
| NDRG4 | 0.84741301 | ENSG00000103034 |
| CDH5 | 1.63143628 | ENSG00000179776 |
| CRABP1 | 2.46242425 | ENSG00000166426 |
| CPXM2 | 1.74617227 | ENSG00000121898 |
| RAI2 | 1.24590163 | ENSG00000131831 |
| ABCA9 | 1.42298851 | ENSG00000154258 |
| SOSTDC1 | 1.18252244 | ENSG00000171243 |
| PCDH1 | 1.59688546 | ENSG00000156453 |
| PI16 | 1.51503289 | ENSG00000164530 |

| | | |
|---|---|---|
| KALRN | 1.4194129 | ENSG00000160145 |
| CPAMD8 | 1.32269839 | ENSG00000160111 |
| U47924.6 | 1.01682406 | ENSG00000240370 |
| AC023137.2 | 1.16743596 | ENSG00000228950 |
| MIR7152 | 3.03645316 | ENSG00000274824 |
| DCHS1 | 1.38392998 | ENSG00000166341 |
| S100B | 1.95057021 | ENSG00000160307 |
| CHST11 | 1.7395662 | ENSG00000171310 |
| CCDC3 | 2.54309434 | ENSG00000151468 |
| CMB9-22P13.1 | 1.11385905 | ENSG00000173727 |
| RAD9A | 0.9880757 | ENSG00000172613 |
| DNM1 | 0.8138897 | ENSG00000106976 |
| SERPINA3 | 2.45165169 | ENSG00000196136 |
| RN7SKP16 | 1.44191178 | ENSG00000222112 |
| CP | 4.04766917 | ENSG00000047457 |
| PRRT2 | 0.86680865 | ENSG00000167371 |
| DTNA | 1.08315879 | ENSG00000134769 |
| AL049758.2 | 1.14936449 | ENSG00000226989 |
| CALCRL | 1.7500092 | ENSG00000064989 |
| C1RL-AS1 | 1.03635173 | ENSG00000205885 |
| PALD1 | 1.21826441 | ENSG00000107719 |
| GPRC5B | 1.48717388 | ENSG00000167191 |
| COL3A1 | 1.87019458 | ENSG00000168542 |
| SEMA3F | 1.16495373 | ENSG00000001617 |
| SEMA6B | 1.11461594 | ENSG00000167680 |
| PIANP | 1.07057472 | ENSG00000139200 |

| | | |
|---|---|---|
| FABP5P7 | 1.00240127 | ENSG00000234964 |
| SLC6A11 | 1.28128009 | ENSG00000132164 |
| GGT5 | 2.39482024 | ENSG00000099998 |
| MIR3134 | 1.04227984 | ENSG00000264354 |
| AC144831.3 | 0.7426091 | ENSG00000274370 |
| OVOS2 | 2.08787498 | ENSG00000177359 |
| SPARCL1 | 2.1381634 | ENSG00000152583 |
| RP11-701H24.8 | 0.58595626 | ENSG00000270246 |
| SOX9 | 0.81808533 | ENSG00000125398 |

† Log2 Fold Change (Peripheral/Central).

**Supplementary Table 5.** List of upregulated and downregulated differentially expressed genes between healthy corneal endothelium and FECD corneal endothelium

| Upregulated in FECD | | | Downregulated in FECD | | |
|---|---|---|---|---|---|
| Gene Symbol | $log_2$FC (F/C) † | ID | Gene Symbol | $log_2$FC (P/C) † | ID |
| DNAJB4 | 0.708890662 | ENSG00000162616 | DNAJA4 | -0.7890435 | ENSG00000140403 |
| MT-TH | 2.050846576 | ENSG00000210176 | RP11-159D12.11 | -1.387151 | ENSG00000278642 |
| CTD-2349P21.9 | 0.916611604 | ENSG00000266490 | CH17-373J23.1 | -2.5920212 | ENSG00000276216 |
| RP11-45M22.2 | 1.01715099 | ENSG00000278864 | SLC47A1 | -1.272223 | ENSG00000142494 |
| GNG4 | 0.941712325 | ENSG00000168243 | HES1 | -1.1847207 | ENSG00000114315 |
| COL4A1 | 1.477525879 | ENSG00000187498 | RNVU1-19 | -3.0533248 | ENSG00000275538 |
| MT-TS2 | 2.342505001 | ENSG00000210184 | KB-1732A1.1 | -1.2704214 | ENSG00000253669 |
| SOX11 | 0.930639587 | ENSG00000176887 | SLC15A4 | -1.2645143 | ENSG00000139370 |
| LBH | 0.89030701 | ENSG00000213626 | ENO1-AS1 | -1.0481405 | ENSG00000230679 |
| SCHIP1 | 1.157485199 | ENSG00000151967 | RNU5D-1 | -1.9178168 | ENSG00000200169 |
| SLC38A1 | 1.044575877 | ENSG00000111371 | SHMT1 | -0.7995588 | ENSG00000176974 |
| MIR6739 | 0.972662428 | ENSG00000277681 | RP11-332H18.3 | -0.886121 | ENSG00000266934 |
| MIR570 | 0.761617814 | ENSG00000207650 | GOLGA2P10 | -0.7620914 | ENSG00000255769 |
| MEST | 1.194596993 | ENSG00000106484 | CPAMD8 | -1.7540899 | ENSG00000160111 |
| SEMA3E | 1.043947499 | ENSG00000170381 | HYKK | -1.0538689 | ENSG00000188266 |
| LOX | 2.11028721 | ENSG00000113083 | LA16c-380F5.1 | -0.9048011 | ENSG00000280062 |
| LAMC1 | 0.653675906 | ENSG00000135862 | IGF2 | -1.2834592 | ENSG00000167244 |
| ETS1 | 0.925413016 | ENSG00000134954 | FAHD2B | -1.0594919 | ENSG00000144199 |
| COL4A2 | 1.521633977 | ENSG00000134871 | RNU11 | -3.0848826 | ENSG00000270103 |
| MIR6728 | 0.765628608 | ENSG00000274258 | C19orf47 | -1.0284236 | ENSG00000160392 |
| RAB11FIP3 | 0.934186715 | ENSG00000090565 | RP11-386I14.4 | -1.7135762 | ENSG00000273338 |

| | | | | | |
|---|---|---|---|---|---|
| RN7SL141P | 0.928111703 | ENSG00000243398 | KCTD10 | -0.6320841 | ENSG00000110906 |
| CACHD1 | 1.079474178 | ENSG00000158966 | RNVU1-6 | -2.6534788 | ENSG00000201558 |
| WTAPP1 | 1.393313509 | ENSG00000255282 | SLCO4A1 | -1.7370058 | ENSG00000101187 |
| RN7SKP137 | 1.207186433 | ENSG00000223006 | GNA14 | -0.6052459 | ENSG00000156049 |
| GUSBP11 | 0.759528228 | ENSG00000228315 | HOMER2 | -0.7431446 | ENSG00000103942 |
| RP11-106M3.2 | 0.957436494 | ENSG00000260729 | MAP2K6 | -0.9961702 | ENSG00000108984 |
| AL513327.1 | 0.755065343 | ENSG00000266239 | LAMC1 | 0.65367591 | ENSG00000135862 |
| IFT122 | 0.59273815 | ENSG00000163913 | APOBEC3G | -0.6759089 | ENSG00000239713 |
| SMURF2 | 0.67643334 | ENSG00000108854 | PLK5 | -1.5062434 | ENSG00000185988 |
| RPL17P50 | 0.606937821 | ENSG00000213700 | TMEM107 | -1.7399177 | ENSG00000179029 |
| MT-TG | 1.549807954 | ENSG00000210164 | DBI | -0.6490784 | ENSG00000155368 |
| RP3-449O17.1 | 0.904069508 | ENSG00000244627 | LSS | -0.704276 | ENSG00000160285 |
| AHR | 0.804011196 | ENSG00000106546 | MYO10 | -0.8633418 | ENSG00000145555 |
| MT-TL2 | 1.857386277 | ENSG00000210191 | MIR616 | -1.5278426 | ENSG00000208028 |
| HS3ST3B1 | 1.500905917 | ENSG00000125430 | U1 | -1.2231046 | ENSG00000278099 |
| MMP14 | 0.647508608 | ENSG00000157227 | APOBEC3C | -0.8042614 | ENSG00000244509 |
| RP11-567P19.1 | 0.943034644 | ENSG00000275927 | RNU1-122P | -1.2869309 | ENSG00000202408 |
| NRG3 | 1.052093619 | ENSG00000185737 | MT-RNR1 | -3.4734977 | ENSG00000211459 |
| SNORD116-2 | 0.896369244 | ENSG00000207001 | PPP1R1B | -2.0577177 | ENSG00000131771 |
| MIR186 | 0.686841656 | ENSG00000207721 | UFSP2 | -0.7397168 | ENSG00000109775 |
| NPHP3-ACAD11 | 1.017998487 | ENSG00000274810 | MFSD6 | -0.9880801 | ENSG00000151690 |
| TENM4 | 0.927840901 | ENSG00000149256 | FAM213B | -0.6018343 | ENSG00000157870 |
| KSR1 | 0.718635851 | ENSG00000141068 | CCND3 | -0.7422697 | ENSG00000112576 |
| Y_RNA | 0.875441899 | ENSG00000200742 | JMJD8 | -0.7343877 | ENSG00000161999 |
| COL26A1 | 0.94237744 | ENSG00000160963 | RNU1-85P | -1.6189343 | ENSG00000200997 |
| RP11-2J18.1 | 0.755200975 | ENSG00000218596 | MIA | -0.993433 | ENSG00000261857 |

| | | | | | |
|---|---|---|---|---|---|
| AC114498.1 | 1.179132706 | ENSG00000276171 | RARA-AS1 | -0.6627013 | ENSG00000265666 |
| CACNA1A | 0.771678598 | ENSG00000141837 | TPD52L1 | -0.6189223 | ENSG00000111907 |
| SMG1 | 0.668579316 | ENSG00000157106 | RP3-453C12.14 | -1.4353494 | ENSG00000275894 |
| RP1-79C4.4 | 1.291814941 | ENSG00000271811 | TRPT1 | -0.6331341 | ENSG00000149743 |
| SLC12A5 | 0.914727073 | ENSG00000124140 | PVALB | -0.8562635 | ENSG00000100362 |
| CCDC80 | 1.120254173 | ENSG00000091986 | HERPUD1 | -1.024772 | ENSG00000051108 |
| DENND3 | 0.618833881 | ENSG00000105339 | MTFP1 | -0.7688482 | ENSG00000242114 |
| MT-TQ | 1.125446676 | ENSG00000210107 | RNU1-120P | -1.806873 | ENSG00000199879 |
| EGFL6 | 1.551866422 | ENSG00000198759 | RPS28P7 | -0.9145977 | ENSG00000227097 |
| RP11-47G11.2 | 1.093125935 | ENSG00000232334 | SERPINA5 | -1.6152077 | ENSG00000188488 |
| GNG12 | 0.6376686 | ENSG00000172380 | RP11-351I21.11 | -1.2126348 | ENSG00000270074 |
| RP11-235E17.6 | 0.761153993 | ENSG00000262903 | RNU5E-4P | -1.9912746 | ENSG00000201801 |
| SNORA66 | 0.66121835 | ENSG00000207523 | NTRK2 | -0.8995629 | ENSG00000148053 |
| SOX4 | 0.795755104 | ENSG00000124766 | AC012370.2 | -0.9692133 | ENSG00000232693 |
| RP11-274H2.5 | 0.706709148 | ENSG00000261051 | PRUNE2 | -0.6883267 | ENSG00000106772 |
| SSH1 | 0.660035811 | ENSG00000084112 | U3 | -2.2155078 | ENSG00000212195 |
| PLEKHG4 | 1.007386259 | ENSG00000196155 | BTG2 | -1.0770596 | ENSG00000159388 |
| GDF5 | 0.961709119 | ENSG00000125965 | SAMM50 | -0.6767797 | ENSG00000100347 |
| MT-TI | 1.188683872 | ENSG00000210100 | EXOSC7 | -0.7407923 | ENSG00000075914 |
| REV3L | 1.02930079 | ENSG00000009413 | U1 | -2.3648121 | ENSG00000206828 |
| IKBIP | 0.788936083 | ENSG00000166130 | PDLIM1 | -0.7771851 | ENSG00000107438 |
| AK5 | 0.925469405 | ENSG00000154027 | RNVU1-15 | -2.7229732 | ENSG00000207205 |
| C1orf198 | 0.611676396 | ENSG00000119280 | ARL 2.00 | -0.7376772 | ENSG00000213465 |
| RP11-138A9.1 | 1.259004792 | ENSG00000271204 | STK35 | -0.9822115 | ENSG00000125834 |
| PLEKHH2 | 0.708507688 | ENSG00000152527 | PNMT | -1.0796392 | ENSG00000141744 |
| PPP1R14BP3 | 0.985607255 | ENSG00000179967 | PPA1 | -0.6523395 | ENSG00000180817 |

| | | | | | |
|---|---|---|---|---|---|
| AL031587.1 | 0.585898666 | ENSG00000280527 | RP11-22N19.2 | -0.8547527 | ENSG00000273320 |
| AGR3 | 0.877684438 | ENSG00000173467 | RNU5A-1 | -1.3001607 | ENSG00000199568 |
| ANTXR1 | 0.794780771 | ENSG00000169604 | RP11-706O15.3 | -1.1317959 | ENSG00000234449 |
| SPSB1 | 0.655207769 | ENSG00000171621 | CATIP-AS1 | -0.7746102 | ENSG00000225062 |
| IQCJ-SCHIP1 | 0.681792597 | ENSG00000250588 | FAM66D | -0.8075462 | ENSG00000255052 |
| MAOA | 0.868570282 | ENSG00000189221 | TAF9 | -0.8855983 | ENSG00000273841 |
| RP11-214O1.2 | 1.025266758 | ENSG00000266709 | PHF7 | -0.8922634 | ENSG00000010318 |
| TTC7A | 0.60477132 | ENSG00000068724 | MT-RNR2 | -2.9195254 | ENSG00000210082 |
| PLOD2 | 0.915578698 | ENSG00000152952 | ALDH1A2 | -1.8307572 | ENSG00000128918 |
| KIAA1468 | 0.869021636 | ENSG00000134444 | CCDC144CP | -1.2379495 | ENSG00000154898 |
| RNU2-2P | 2.523824322 | ENSG00000222328 | AKR1B1 | -0.9087979 | ENSG00000085662 |
| USP40 | 0.618352953 | ENSG00000085982 | WDR61 | -0.8586815 | ENSG00000140395 |
| SHB | 0.598347174 | ENSG00000107338 | ST6GALNAC2 | -0.9902373 | ENSG00000070731 |
| PANX2 | 0.876811249 | ENSG00000073150 | ZNF839 | -1.1108057 | ENSG00000022976 |
| U3 | 0.785740138 | ENSG00000238297 | AC010761.14 | -0.7647538 | ENSG00000267729 |
| C5orf42 | 0.686180934 | ENSG00000197603 | CBR1 | -0.782356 | ENSG00000159228 |
| CLK4 | 0.811414026 | ENSG00000113240 | RANBP3L | -1.3195762 | ENSG00000164188 |
| RP11-274H2.3 | 0.753828259 | ENSG00000240032 | H19 | -1.5372582 | ENSG00000130600 |
| HPGD | 1.353149389 | ENSG00000164120 | TOB1-AS1 | -0.7425564 | ENSG00000229980 |
| BCORL1 | 0.629716193 | ENSG00000085185 | RNU2-63P | -1.769004 | ENSG00000222724 |
| SKIL | 0.615019194 | ENSG00000136603 | TNNT3 | -1.3302878 | ENSG00000130595 |
| RRP7BP | 0.634522357 | ENSG00000182841 | ETNPPL | -1.3755149 | ENSG00000164089 |
| ARNTL | 0.905899493 | ENSG00000133794 | MYCL | -1.0628845 | ENSG00000116990 |
| AC025279.1 | 0.797101416 | ENSG00000278509 | TMOD1 | -1.5540138 | ENSG00000136842 |
| MT-TN | 1.047325971 | ENSG00000210135 | H1F0 | -0.7579533 | ENSG00000189060 |
| MIR6888 | 0.592629733 | ENSG00000275141 | LINC01220 | -0.8679497 | ENSG00000259687 |

| | | | | | |
|---|---|---|---|---|---|
| GALNT18 | 0.736115484 | ENSG00000110328 | ARF4-AS1 | -0.8234157 | ENSG00000272146 |
| LOXL2 | 1.118400719 | ENSG00000134013 | CTD-2659N19.9 | -0.8215062 | ENSG00000267212 |
| RNU7-124P | 0.800503445 | ENSG00000251745 | MIR4755 | -0.8570619 | ENSG00000264616 |
| SERPINH1 | 0.645679121 | ENSG00000149257 | BOLA3 | -0.6316541 | ENSG00000163170 |
| SEMA5A | 0.763761536 | ENSG00000112902 | NATD1 | -0.8763041 | ENSG00000274180 |
| HS3ST3A1 | 0.979435586 | ENSG00000153976 | PSMD5-AS1 | -0.9653725 | ENSG00000226752 |
| FBN1 | 1.249202804 | ENSG00000166147 | MIR5188 | -1.2550662 | ENSG00000265345 |
| DKK 2.00 | 0.706985928 | ENSG00000155011 | LYNX1 | -0.7152332 | ENSG00000180155 |
| ZFP36L2 | 0.606525326 | ENSG00000152518 | BAIAP2 | -0.6863176 | ENSG00000175866 |
| RP11-430B1.1 | 0.863556659 | ENSG00000259398 | RNVU1-14 | -2.294969 | ENSG00000207501 |
| SERPINE2 | 1.199629428 | ENSG00000135919 | DHCR24 | -0.6634773 | ENSG00000116133 |
| SMAD6 | 0.720666847 | ENSG00000137834 | ZNF226 | -1.3547641 | ENSG00000167380 |
| AC008740.1 | 1.141509313 | ENSG00000276555 | RNU2-1 | -1.2493647 | ENSG00000274585 |
| ODC1 | 0.679310594 | ENSG00000115758 | NAA38 | -0.6357854 | ENSG00000183011 |
| MIR1231 | 0.901958358 | ENSG00000221028 | WARS2 | -0.6029931 | ENSG00000116874 |
| RGS11 | 0.820502395 | ENSG00000076344 | MVK | -1.1563734 | ENSG00000110921 |
| APELA | 0.766631527 | ENSG00000248329 | MPND | -0.6768035 | ENSG00000008382 |
| SCARNA20 | 0.710556573 | ENSG00000252577 | RP1-193H18.2 | -0.6869185 | ENSG00000267194 |
| FEZ1 | 0.85786979 | ENSG00000149557 | RNVU1-4 | -1.9892773 | ENSG00000277610 |
| ZMYM6NB | 0.800699821 | ENSG00000243749 | PGR | -1.0384737 | ENSG00000082175 |
| HNRNPU-AS1 | 0.986755424 | ENSG00000188206 | TUBA4A | -0.6144779 | ENSG00000127824 |
| GAS1 | 0.656979172 | ENSG00000180447 | KLHDC4 | -0.8762203 | ENSG00000104731 |
| KIF5C | 0.682853836 | ENSG00000168280 | MCCC1 | -0.6408355 | ENSG00000078070 |
| POLR2J2 | 0.651816911 | ENSG00000228049 | DPH5 | -0.7737018 | ENSG00000117543 |
| FZD1 | 0.688713799 | ENSG00000157240 | WASH7P | -0.7368479 | ENSG00000226210 |
| PMEPA1 | 0.73479321 | ENSG00000124225 | RP11-727F15.12 | -0.7610382 | ENSG00000269176 |

| | | | | | |
|---|---|---|---|---|---|
| TPM4 | 0.667626945 | ENSG00000167460 | CLRN1 | -1.0267051 | ENSG00000163646 |
| FOXP4 | 0.861868102 | ENSG00000137166 | U1 | -2.4139174 | ENSG00000274210 |
| IGFBP7 | 1.20161903 | ENSG00000163453 | STRA13 | -0.6434553 | ENSG00000169689 |
| RP11-355O1.11 | 0.875816854 | ENSG00000261094 | AMIGO2 | -0.8012739 | ENSG00000139211 |
| | | | AP000322.53 | -0.9658449 | ENSG00000243627 |
| | | | FRMD3 | -0.9396735 | ENSG00000172159 |
| | | | ANAPC15 | -0.7659732 | ENSG00000110200 |
| | | | NIPSNAP3B | -0.6205356 | ENSG00000165028 |
| | | | SNHG16 | -0.8240588 | ENSG00000163597 |
| | | | MPP6 | -0.9736188 | ENSG00000105926 |
| | | | ACY1 | -0.7740815 | ENSG00000243989 |
| | | | MFSD11 | -0.67444 | ENSG00000092931 |
| | | | CKMT1A | -0.8892985 | ENSG00000223572 |
| | | | CH507-396I9.3 | -0.9481748 | ENSG00000278961 |
| | | | HLF | -0.9394276 | ENSG00000108924 |
| | | | COTL1 | -0.7864635 | ENSG00000103187 |
| | | | TBC1D2 | -0.7425638 | ENSG00000095383 |
| | | | S100A6 | -0.7154376 | ENSG00000197956 |
| | | | FRMPD1 | -0.768885 | ENSG00000070601 |
| | | | CTB-58E17.1 | -0.637795 | ENSG00000277969 |
| | | | ATMIN | -0.6041622 | ENSG00000166454 |
| | | | U1 | -0.8509197 | ENSG00000274428 |
| | | | RNU12 | -1.9321779 | ENSG00000270022 |
| | | | ALDH3A1 | -2.0980491 | ENSG00000108602 |
| | | | RP11-318A15.8 | -0.9140657 | ENSG00000277382 |
| | | | AC079922.3 | -0.6347921 | ENSG00000237753 |

| | | | |
|---|---|---|---|
| | CKMT1B | -0.9115309 | ENSG00000237289 |
| | U2 | -1.247066 | ENSG00000276596 |
| | TEF | -0.7153274 | ENSG00000167074 |
| | ACAD8 | -0.7189112 | ENSG00000151498 |
| | U2 | -1.2156278 | ENSG00000277903 |
| | U2 | -1.3065945 | ENSG00000275616 |
| | PCYT2 | -0.6027845 | ENSG00000185813 |
| | KRT15 | -1.45751 | ENSG00000171346 |
| | ETV4 | -0.9550436 | ENSG00000175832 |
| | OSGIN1 | -0.8553094 | ENSG00000140961 |
| | U2 | -1.2638955 | ENSG00000278774 |
| | OGDHL | -0.6253224 | ENSG00000197444 |
| | REEP6 | -0.9245455 | ENSG00000115255 |
| | RP11-467P9.1 | -0.7496419 | ENSG00000272735 |
| | ZNF775 | -0.9070976 | ENSG00000196456 |
| | FRMD4A | -0.8264057 | ENSG00000151474 |
| | CLEC4GP1 | -1.1434544 | ENSG00000268297 |
| | DAPL1 | -1.1397829 | ENSG00000163331 |
| | RP11-229P13.25 | -0.7973382 | ENSG00000260190 |
| | U2 | -1.2360162 | ENSG00000274862 |
| | CTB-25B13.12 | -0.6589847 | ENSG00000267317 |
| | BCKDHA | -0.6000996 | ENSG00000248098 |
| | U2 | -1.2517245 | ENSG00000275219 |
| | MSMP | -1.7243508 | ENSG00000215183 |
| | GJB6 | -1.6455922 | ENSG00000121742 |
| | MPP1 | -0.9311646 | ENSG00000130830 |

| | | |
|---|---|---|
| KRT3 | -2.1743355 | ENSG00000186442 |
| EGR1 | -1.8276176 | ENSG00000120738 |
| KRT12 | -3.3025033 | ENSG00000187242 |
| AC245033.1 | -0.7901745 | ENSG00000221095 |
| TOMM40L | -0.6335148 | ENSG00000158882 |
| U2 | -1.2324611 | ENSG00000274062 |
| RPL18AP3 | -0.6076384 | ENSG00000213442 |
| U2 | -1.1431185 | ENSG00000274452 |
| CLDN10 | -1.0355649 | ENSG00000134873 |
| U2 | -1.3160064 | ENSG00000278048 |
| U2 | -1.2255814 | ENSG00000278591 |
| SYT13 | -0.9656074 | ENSG00000019505 |
| MYZAP | -0.9343713 | ENSG00000263155 |
| USP32P3 | -0.9658377 | ENSG00000189423 |
| VAV3 | -0.8755892 | ENSG00000134215 |
| COL17A1 | -1.8498714 | ENSG00000065618 |
| ZNF789 | -0.7119071 | ENSG00000198556 |
| U1 | -1.5808566 | ENSG00000270722 |
| TSC22D3 | -1.1102007 | ENSG00000157514 |
| GJB2 | -1.8822585 | ENSG00000165474 |
| C11orf73 | -0.7773328 | ENSG00000149196 |
| ST3GAL5-AS1 | -0.7147769 | ENSG00000232504 |
| NOL12 | -0.7663189 | ENSG00000273899 |
| DHCR7 | -0.6968409 | ENSG00000172893 |
| CITF22-49E9.3 | -0.8211124 | ENSG00000278869 |
| RORC | -1.0422719 | ENSG00000143365 |

| | | | |
|---|---|---|---|
| | FAM106CP | -0.6243244 | ENSG00000266486 |
| | RNF39 | -0.8755947 | ENSG00000204618 |
| | HIF3A | -0.8599833 | ENSG00000124440 |
| | THUMPD3-AS1 | -0.6270871 | ENSG00000206573 |
| | SDHAP2 | -0.8202056 | ENSG00000215837 |
| | TMEM189-UBE2V1 | -0.9130345 | ENSG00000124208 |
| | U2 | -1.2230469 | ENSG00000274432 |
| | GRB10 | -0.8775491 | ENSG00000106070 |
| | ITGB3BP | -0.80622 | ENSG00000142856 |
| | CYYR1 | -0.6449731 | ENSG00000166265 |
| | GHITM | -0.638003 | ENSG00000165678 |
| | DBNL | -0.6009732 | ENSG00000136279 |
| | MIR3193 | -0.8377913 | ENSG00000264395 |
| | AGPAT3 | -0.839273 | ENSG00000160216 |
| | ETFB | -0.6274966 | ENSG00000105379 |
| | WDR74 | -1.0088433 | ENSG00000133316 |
| | EPHB1 | -0.780306 | ENSG00000154928 |
| | GCOM1 | -0.7772417 | ENSG00000137878 |
| | SKP2 | -0.8504147 | ENSG00000145604 |
| | RBP1 | -1.024938 | ENSG00000114115 |
| | FA2H | -0.7959968 | ENSG00000103089 |
| | CNTD2 | -0.9861462 | ENSG00000105219 |
| | U2 | -1.2948188 | ENSG00000273709 |
| | AC005702.1 | -0.7936256 | ENSG00000263422 |
| | HSPB7 | -0.869861 | ENSG00000173641 |
| | KLF10 | -1.2762091 | ENSG00000155090 |

| | | | |
|---|---|---|---|
| | HGH1 | -0.6301285 | ENSG00000235173 |
| | ADAMTS7P4 | -1.0742699 | ENSG00000218052 |
| | USP45 | -0.8687223 | ENSG00000123552 |
| | RBP7 | -1.0403233 | ENSG00000162444 |
| | PTGDS | -0.7634871 | ENSG00000107317 |
| | RP11-415J8.3 | -0.7676736 | ENSG00000225313 |
| | ARHGAP40 | -1.210462 | ENSG00000124143 |
| | MIR5004 | -0.8303262 | ENSG00000264085 |
| | RNA5SP146 | -0.7754518 | ENSG00000222675 |

† Log2 Fold Change (FECD/Healthy Control)
